# Resolving the conflict between antibiotic production and rapid growth by recognition of peptidoglycan of susceptible competitors

**DOI:** 10.1101/2021.02.07.430110

**Authors:** Harsh Maan, Jonathan Friedman, Ilana Kolodkin-Gal

## Abstract

Microbial communities employ a variety of complex strategies to compete successfully against competitors sharing their niche, with antibiotic production being a common strategy of aggression. Here, by systematic evaluation of all non-ribosomal peptides (NRP) produced by *B. subtilis* clade, we revealed that they acted either synergistically or additively to effectively eliminate phylogenetically distinct competitors. All four major NRP biosynthetic clusters were also imperative for the survival of *B. subtilis* in a complex community extracted from the rhizosphere. The production of NRP came with a fitness cost manifested in growth inhibition, rendering NRP synthesis uneconomical when growing in proximity to a phylogenetically close species, carrying resistance against the same antibiotics. To resolve this conflict and ease the fitness cost, NRP production was only induced by the presence of peptidoglycan cue from a sensitive competitor. These results experimentally demonstrate a general ecological concept – closely related communities (“self”) are favoured during competition, due to compatibility in attack and defence mechanisms.

## Introduction

In terrestrial microenvironments, bacteria are present predominantly in architecturally complex and specialised multispecies communities ^1^. In many instances, these communities provide beneficial effects to other organisms, e.g., biocontrol agents form biofilms on the surface of plant roots, thereby preventing the growth of bacterial and fungal pathogens ^2^. Soil and plant-associated microbial populations survive in this extremely competitive niche by producing a broad arsenal of antibiotics and evolving complex antibiotic-resistance mechanisms ^3,4^. Social structure was shown to mediate competition between populations, suggesting a role for antibiosis in shaping cohesive communities ^5^.

*Bacillus subtilis* clade contains several species of Gram-positive, soil dwelling, beneficial bacteria, that employ multiple strategies to compete with neighbour communities ^2,6–12^.One major class of antibiotics produced by bacteria are the Non-Ribosomal Peptides (NRP), which are synthesized by large multi-enzyme complexes of NRP synthases (NRPs) ^2,13,14^. *B. subtilis* encodes four different NRP biosynthetic clusters, which are responsible for the biosynthesis of surfactin, bacillaene, bacilysin and plipastatin ^2,15–18^.

The most well-characterized NRP is the cyclic lipopeptide surfactin ^19,20 21^. Surfactin is a small cyclic lipopeptide induced during the development of genetic competence ^22^. The machinery for surfactin synthesis is encoded within the *srfAA-AB-AC-AD* operon ^23^. Surfactin is a powerful surfactant with antibacterial ^24^ and antifungal properties ^25^ and is composed of an amphipathic, cyclic heptapeptide head group that is interlinked with a hydrophobic β-hydroxy fatty acid tail, comprising 12–16 carbon atoms ^26–28^. These features enable the surfactin molecule to interact with, and disrupt the integrity of, cellular membranes ^29^.

Bacillaene and dihydrobacillaene ^15,30^, are linear antimicrobial macrolides with two amide bonds and are synthesized by the PksC-R cluster mega-complex, which is composed of 13 PKS and three NRPS modules ^30^. The expression of *pks* genes requires the master regulator for biofilm formation, Spo0A, and promotes the competitiveness of biofilm communities ^31^.

Bacilysin is a non-ribosomal dipeptide is composed of the amino acids L-alanine and L-anticapsin, and acts as an antibiotic with activities against a wide range of bacterial and fungal pathogens ^32^. Its synthesis is controlled mainly by the *bac* operon *(bacABCDE)* and is regulated by other enzymes such as thymidylate synthase, homocysteine methyl transferase and the oligopeptide permeases ^33^. This dipeptide undergoes peptidase-mediated proteolysis to release L-anticapsin, which is a competitive inhibitor of glucosamine synthase ^32^. This dipeptide can also be produced as chlorotetain, with an unusual chlorine-containing amino acid ^34,35^.

Lastly, plipstatin is synthesized non-ribosomally by five fengycin synthetases (*ppsA-E*), that are regulated by DegQ, a master regulator of the transition from a motile cell state to a biofilm-forming state ^36^ DegQ is activated by catabolite repression and is under the control of the DegSU two-component system, that control the formation of the extracellular matrix, sporulation, protease production and motility ^37,38^. ComA is a response regulator acting in the two-component regulatory system ComP/ComA involved in genetic competence and surfactin production ^39^ It was recently reported that ComA influences the transcription of more than 100 genes ^40^, including the genes involved in the biosynthesis of Bacillaene ^31^ and Bacilysin ^41^.*B. subtilis* strains were previously shown to distinguish related (Kin) and non-related species during swarming^9,10^. The activation of the cell wall stress response was demonstrated during non-kin interactions; however, no single deletion or a combination of deletions had a detectable effect on non-kin boundaries ^9^. Although the production of antimicrobials were suggested to play a role in the boundaries between non-kin swarms ^9^, the role of NRP clusters in non-kin recognition was not directly examined.

The production of several potent antibiotic clusters in *B. subtilis*, together with its capacity to distinguish non-kin *B. subtilis* strains allows us to utilize a genetic approach to investigate fundamental questions regarding antibiosis. For example, what is the fitness benefit of carrying several antibiotics biosynthetic clusters within a single genome? Is there a trade-off between antibiotic biosynthesis and growth? How can bacteria avoid producing costly antibiotics against members of the community that are immune to their action?

To resolve these open questions, we systematically explored the roles of NRPs antibiotics biosynthetic clusters and the transcription from their promoters during interactions between related *bacillus* species.

We found that *bacilli* species resolve the conflict between NRP production and growth by increasing the transcription of NRP only when competing against phylogenetically distant species, which are sensitive to this class of antibiotics. In contrast, NRP production was not enhanced by the presence of NRP-resistant members of the genus. Purified non-self peptidoglycan (PG), a polymer constituting the outmost layer of the cell wall of the competitors, was sufficient to induce the expression of antibiotic biosynthesis promoters, while self-produced peptidoglycan was either inert or inhibitory to their expression. Furthermore, a single transcription factor, ComA, was essential to actively distinguish non-self PG from self-PG. These results demonstrate how the formation of communities composed of closely related species is favoured during competition, due to compatibility in antibiotic production and sensitivity. Furthermore, they suggest a new mechanism by which microbial populations resolve the conflict between antibiotic production and growth.

## Results

### *B. subtilis* Eliminates Non-self Competitors by NRP Antibiotics

To determine how NRP biosynthetic clusters interact during microbial competition, we first examined the competition between *B. subtilis* and several representative *bacilli* species spread over various phylogenetic distances (Figure 1A). We performed the experiment on a biofilm medium B4 ^42^, as it was previously demonstrated that antibiotic production is frequently induced together with biofilm formation in the *Bacillus* clade ^43–46^.

**Figure 1:**
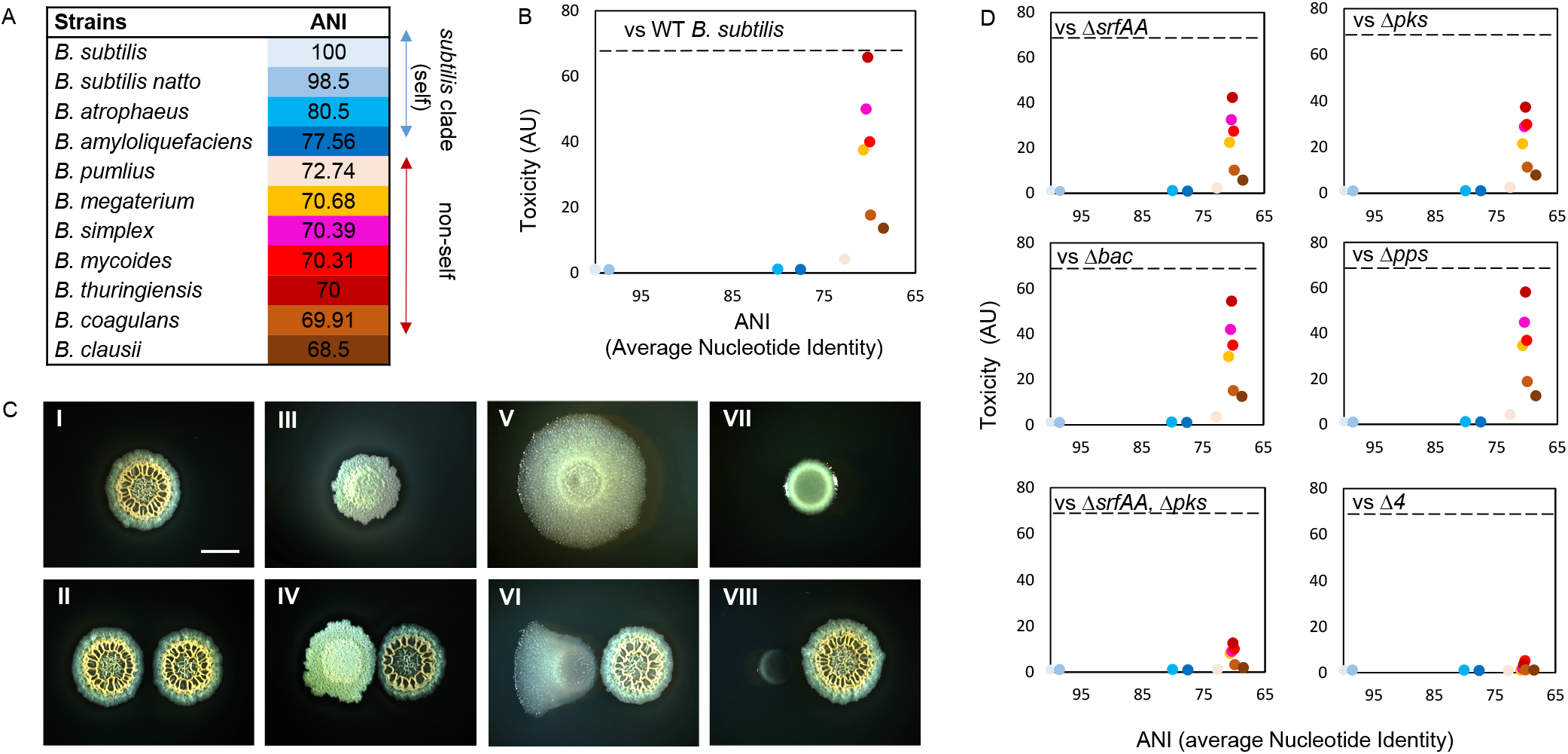
The contribution of non-ribosomal peptides to within genus competition. **A)** Average nucleotide identity of indicated *bacillus* species compared with *B. subtilis* **B)** Toxicity of *B. subtilis* wild-type strain towards competitors. Toxicity were determined as the [growth alone] / [growth in competition against *B. subtilis]* (AU): Biofilm cells were harvested and colony forming units (CFU) were calculated alone, and during co-inoculation. X-axis represents the average nucleotide identity between indicated *Bacillus species*. Dashed Line: The toxicity of the WT *B. subtilis* **C)** Colonies were imaged 48 hrs post inoculation **I)** *B. subtilis* colony grown alone **II)** in competition against *B. subtilis* **III)** *B. amyloliquefaciens* grown alone **V)** *B. thuringiensis* grown alone **VI)** in competition against *B. subtilis* **VII)** *B. coagulans* grown alone **VIII)** in competition against *B. subtilis*. Scale bar = 1mm **D)** The toxicity of *B. subtilis* single mutants, a double mutant for *srfAA* and *pks*, and a quadruple mutant for all NRPs operons *srfAA-srfAD, pksC-pksR, bacA-ywfG and ppsA –ppsE* (Δ*4*) towards competitors was evaluated as in B.

Phylogenetic distance between *B. subtilis* and its competitors was quantified using the nucleotide level genomic similarity between the coding regions of two genomes, expressed as ANI (Average Nucleotide Identity) ^47^ A competition assay was developed to test the potential interactions between *B. subtilis* and its competitors. The degree of toxicity exhibited by *B. subtilis* towards competing *bacilli* was calculated by dividing the culturable cell counts (as judged by colony forming units, CFU) of biofilms grown in isolation by CFU of biofilms competing against *B. subtilis*.

Our results showed that members of the *subtilis* clade (“self”) – *B. subtilis 3610, B. subtilis natto, B. amyloliquefaciens*, and *B. atropheus* – coexisted well. In contrast, phylogenetically distinct *species* (“non-self”) – *B. megaterium, B. mycoides, B. simplex, B. thuringiensis, B. coagulans and B. clausii* – were eliminated by *B. subtilis*. The maximum degree of toxicity exhibited by *B. subtilis* against *non-self-bacilli* was ≃ 65 folds (Figure 1B). Results were consistent with alterations in colony morphologies and growth during interactions: members of the “non-self” group struggled to grow in proximity to *B. subtilis*, while members of the same clade were largely unaffected by this interaction (Figures 1B and C, Figure S1). For all interactions, *B*. subtilis NCIB 3610 itself was largely unaffected (Figure S2)

We studied the contribution of four different NRPs gene clusters to the antibiosis: *srfAA-srfAD, pksC-pksR, bacA-ywfG and ppsA-ppsE;* responsible for the biosynthesis of surfactin, bacillaene, bacilysin and plipastatin, respectively. The degree of toxicity exhibited by WT (wild type) *B. subtilis* (maximum 65 folds) was used as a reference. When *bacilli* were competed against a mutant lacking either *srfAA* (Δ*srfAA*) or *pksC-pksR* (Δ*pks*), the toxicity was reduced to 37 and 42-fold respectively (Figure 1D). In contrast, the toxicity of strains lacking *bacA*-Δ*ywfG* (Δ*bac*) *and ppsA-ppsE* (Δ*pps*) was similar to that of the WT strain (Figure 1D). Next, we examined combinations of double deletions. Only a double mutant lacking surfactin and bacillaene biosynthesis (Δ*srfAA*, Δ*pks*) was clearly compromised in its toxicity compared to either of the single deletion mutants (Figure 1D and S3). Furthermore, a quadruple mutant lacking all NRP biosynthetic clusters (Δ*4*) lost all toxicity towards “non-self” genus members (Figure 1D).

### Secreted Products of NRP Biosynthetic Clusters Interact Additively and Synergistically

Consistent with the notion that the toxicity if mediated by a secreted antibiotic product, conditioned medium from WT *B. subtilis* NRP antibiotics was toxic to “non-self” *bacilli*, but the toxicity of conditioned medium produced by the different NRPS deletion mutants was compromised (Figures 2A and S4). Similarly to their contribution during interspecies interaction on plate, conditioned media containing surfactin and bacillaene contributed most of the toxicity towards “non-self” community members, while maintaining minimal or no toxicity towards “self” (Figure 2A). Overall, these results indicate that the “self” members of *B. subtilis* clade were largely resistant to NRP antibiotics, either alone or in combination, while the competitors outside of the clade were sensitive to the combinatory activity of these clusters.

**Figure 2:**
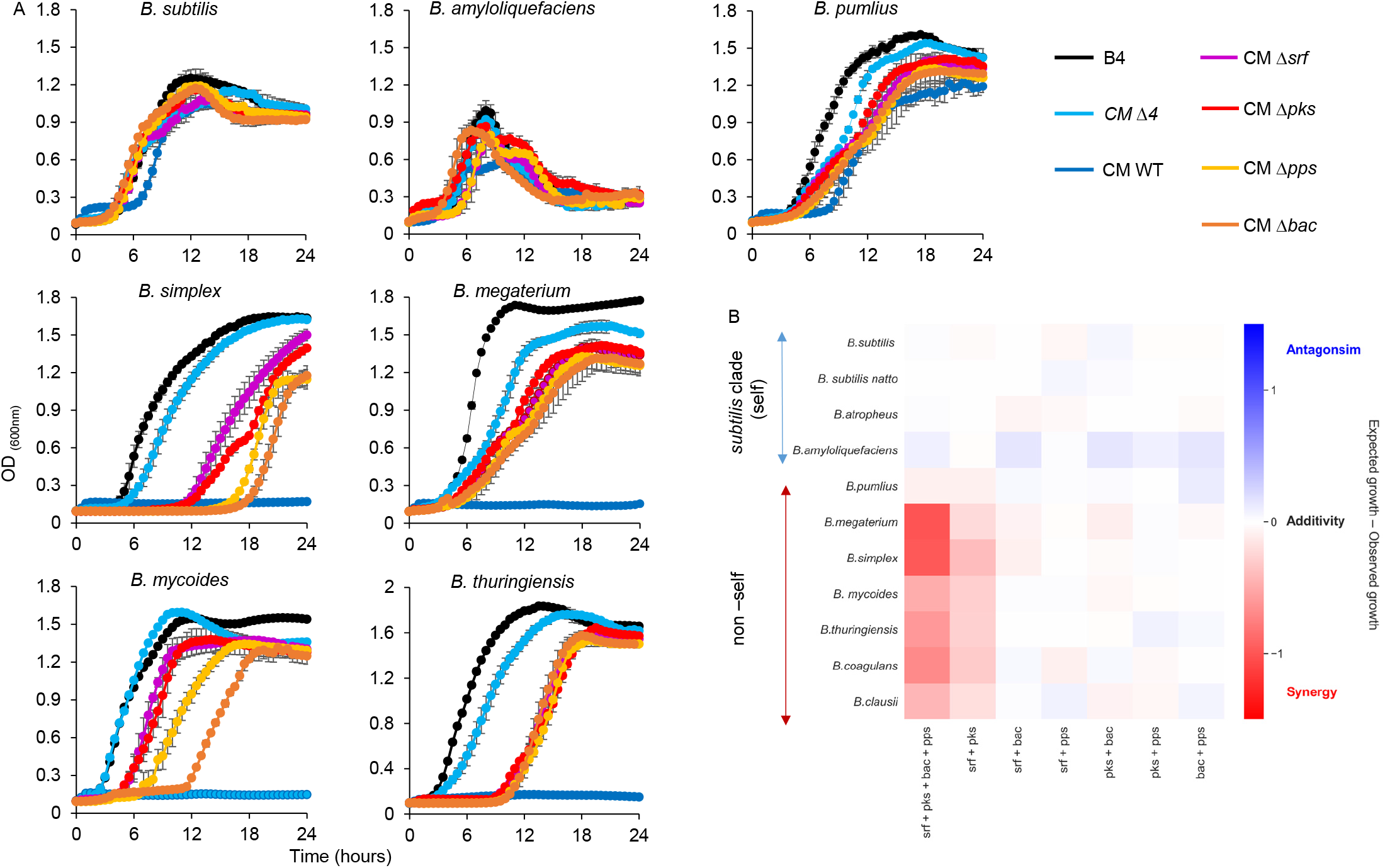
The Interaction between Secreted NRPs Products. **A)** Planktonic growth of the indicated species was monitored either in B4 medium or B4 medium supplemented with CM of the parental strain (WT) or its indicated NRPs biosynthetic clusters mutants. **B)** Heat map showing the calculated interactions between NRP antibiotics. Interaction score = 0 represents no interaction, <0 represents antagonistic interaction and > 0 represents synergistic interactions. Among all the possible NRP combinations, surfactin (*srf*) and bacillaene (*pks*) acted synergistically against non-self-members.

Multiple agents synergize if they are more effective in combination than one would expect by adding their individual effects. To determine if NRP biosynthetic cluster interact, their observed combined effect was compared with a reference that predicts how individual effects would add up independently. Relying on the genetic analysis of antibiosis described in Figure 1, we compared the *observed* combined toxicity of multiple antibiotics to the one *expected* from the Bliss additivity model ^19^

Interactions score of 0 indicated additivity (antibiotics that act independently), scores greater than 0 represented antagonism and scores less than 0 indicated synergy. No significant antagonism was identified in any of the antibiotic combinations studied. In addition, none of the pairs showed any synergy against “self”. In contrast, the pair bacillaene – surfactin acted synergistically against “non-self” *Bacilli*, while the rest of the pairs showed additive interactions. When the effects of all four antibiotics were combined, synergism against “non-self” increased up to 5-fold (Figure 2B).

We then asked whether these antibiotics act similarly in highly complex communities. Native rhizosphere was extracted from the roots and adhering soil of annual plants organic *Solanum lycopersicum* (Figure 3A), and competed versus the parental strain and its mutants.

**Figure 3:**
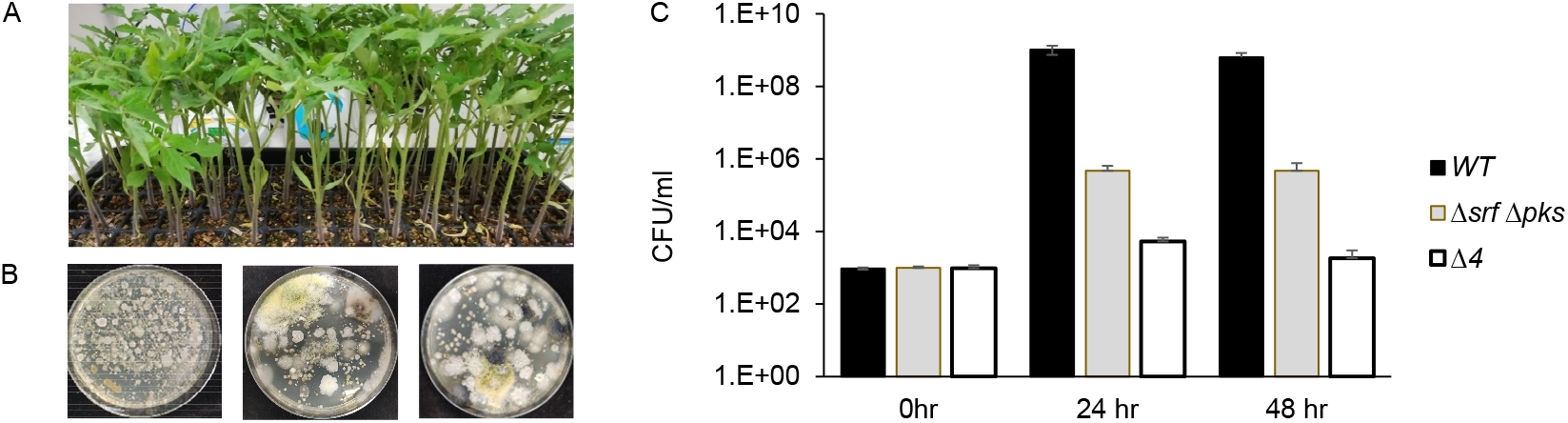
All Four NRPS Products Are Required to Compete in Complex Community. **A)** The tomato plants used for rhizosphere extraction. **B)** The diversity of rhizosphere community microbiome on LB plates. **C)** The abundance of WT, Δ*srfAA*Δ*pks* and Δ*4* at different time points after being introduced into the extracted rhizosphere community. Results are average and SD of three independent experiments.

Specifically, we compared the relative contribution of NRPs cluster by a competition essay: the extracted rhizosphere communities (Figure 3A and B) were competed against equal amounts of the parental strain, a double mutant for bacillaene and surfactin and a quadruple mutant for all major NRPs clusters (Δ*4*). At T=0 and prior to interacting with the complex community, the quantity of all strains were compatible (Figure 3B). Within 24 hours within the complex community, the parental strain gained advantage over the double mutant (Δ*srfAA*, Δ*pks*) and the quadruple mutant abundance was significantly decreased compared with both parental strain and a double mutant (Figure 3C). These results support the accumulative contribution of all NRP biosynthetic clusters to microbial competition observed by us *in vitro* (Figures 1 and 2).

### NRP Biosynthetic Cluster Expression is Induced by Sensitive Competitors to Reduce the Burden on Microbial Growth

NRP gene clusters encode large complexes composed of many protein subunits. Considering the burden of their production, we wondered whether these operons are differentially transcribed during interaction with the resistant “self” and the sensitive “non-self” competitors.

Using flow cytometry, we examined the expression of transcriptional reporters P*_srfAA_-yfp*, P*_pksC_*-*mKate*, P*_bacA_*-*gfp* and P*_ppsA_*-*gfp* in biofilm colonies. NRP producing cells consisted distinct subpopulations (Figure S5), indicating that these antibiotics act as a common good and are shared between biofilm community members. We then asked whether interspecies competition would have an impact on their expression. The number of cells expressing each NRP and their respective mean intensity of fluorescence remained constant or even decreased while interacting with “self” *subtilis* clade members. On the other hand, both increased during competition against “non-self” *bacilli* (Figure 4A), and Table S4), linking transcriptional activity of these promoters and the potential fitness benefits of NRP production.

**Figure 4:**
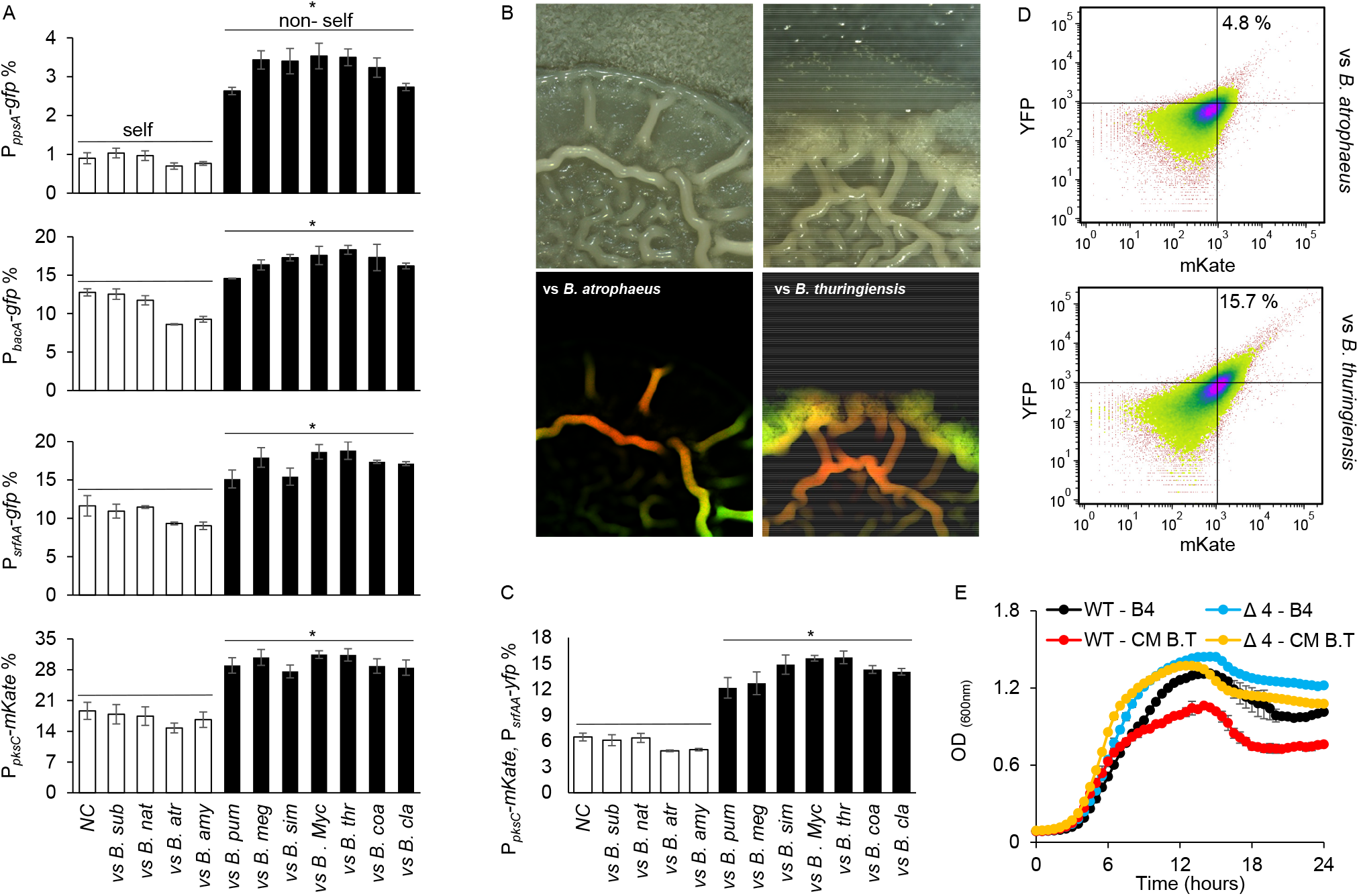
Distinguishing Self and Non-Self Regulates the Transcription from the Promoters of NRP Biosynthetic Clusters. **A)** Indicated reporter strains were analyzed either during growth either in isolation (NC) or during indicated interactions using flow cytometry. Data were collected from 24 h post inoculation. Y-axis represents fluorescent populations (100,000 cells were counted). Experiments were performed in triplicates, a representative example is shown from three independent experiments. * (p < 0.001) between self and non-self *bacillus species*. **B)** A dual reporter strain carrying both P*_srfAA_*-*yfp* (surfactin) and P*_pks_*-*mKate* (bacillaene) was grown as in Figure 1 and imaged using Zeiss stereomicroscope at 44-x magnification. Upper panel (Phase), Bottom panel (Fluorescence). An increase in the fluorescence of dual population was observed when competed against non-self-bacilli – *B. thuringiensis*. **C)** A dual reporter strain carrying both P*_srfAA_*-*yfp* and P*_pks_*-*mKate* was analyzed either during growth either in isolation (NC) or during indicated interactions using flow cytometry. Data were collected from 24 h post inoculation. Y-axis represents fluorescent populations (100,000 cells were counted). Experiments were performed in triplicates, a representative example is shown from three independent experiments. (p < 0.001) between self and non-self *bacillus species* **D)** Quantification of dual reporter carrying both P*_srfAA_*-*yfp* and P*_pks_*-*mKate*. Top upper right panel represents one subpopulation of cells. This subpopulation is detected in the YFP and mKate channel because it expresses the two reporters simultaneously. An increase in the dually expressing population was observed during competition against *B. thuringiensis*. Data were collected from 24 h post inoculation. Y-axis represents YFP fluorescence channel and x - axis represents mKate fluorescence channel (100,000 cells were counted). **E)** Growth of indicated strains applied with conditioned medium from *B. thuringiensis*. Reduction of growth by the *B. thuringiensis* supernatant was depend on the presence of NRP biosynthetic clusters

An additional layer of regulation on NRPs production could be the presence within the community of subpopulations specializing on expression of one or more biosynthetic clusters. T o further study the division of labour at single cell level, a dual transcriptional reporter, harbouring P*_srfAA_*-*yfp*, P*_pksC_*-*mKate*, was constructed and the expression of these alleles was assessed simultaneously during growth in isolation, and during competition. A distinct population expressing from both surfactin and bacillaene promoters was detected (Figure 4B). This further suggests that NRP-expressing subpopulations of *B. subtilis*, exhibit, to some extent, differentiation into specialized subpopulations. Furthermore, we observed a two-fold increase in the population expressing both gene clusters in the presence of “non-self” *bacilli*, which are sensitive to those antibiotics (Figure 4C).

In order to understand whether the enhanced production of NRP affects the growth of *B. subtilis*, we monitored growth rates of WT *B. subtilis* and its NRP deletion mutants. Indeed, the growth of parental strain was significantly lower than that of all mutants tested – and especially as compared to that of the quadruple mutant (Figure S6). As we observed that *B. subtilis* increased the transcription of NRPs gene clusters when competing against “non-self” *bacilli*, we tested whether this induction increased the burden of NRPs synthesis on the growth of *B. subtilis*. The *B. subtilis* parental strain and quadruple mutant were grown with or without the conditioned medium from the “self” and “non-self” *bacilli*. The inducing conditioned medium from “non-self” *bacilli* significantly increased the growth inhibition of the parental strain, but not of the quadruple mutant (Figure 4D), indicating that the induction of NRP synthesis by non-self has deleterious effects on the producers.

### Peptidoglycan Recognition acts upstream to ComA to Induce Non-Self Recognition and Transcription from NRP promoters

We asked whether NRPs induction by specifically sensing a secreted signal from “non-self”. For real-time measurements of gene expression, we used *B. subtilis* strains harbouring the luciferase reporters P*_pksC_*-*lux* and P*_srfAA_*-*lux*. As shown (Figure 5A and Figure S7), conditioned medium containing physiological concentrations of secreted molecules of “non-self” competitors was sufficient to induce the transcription of both surfactin and bacillaene. In contrast, the secretions of resistant “self” competitors failed to induce surfactin and bacillaene expression.

**Figure 5:**
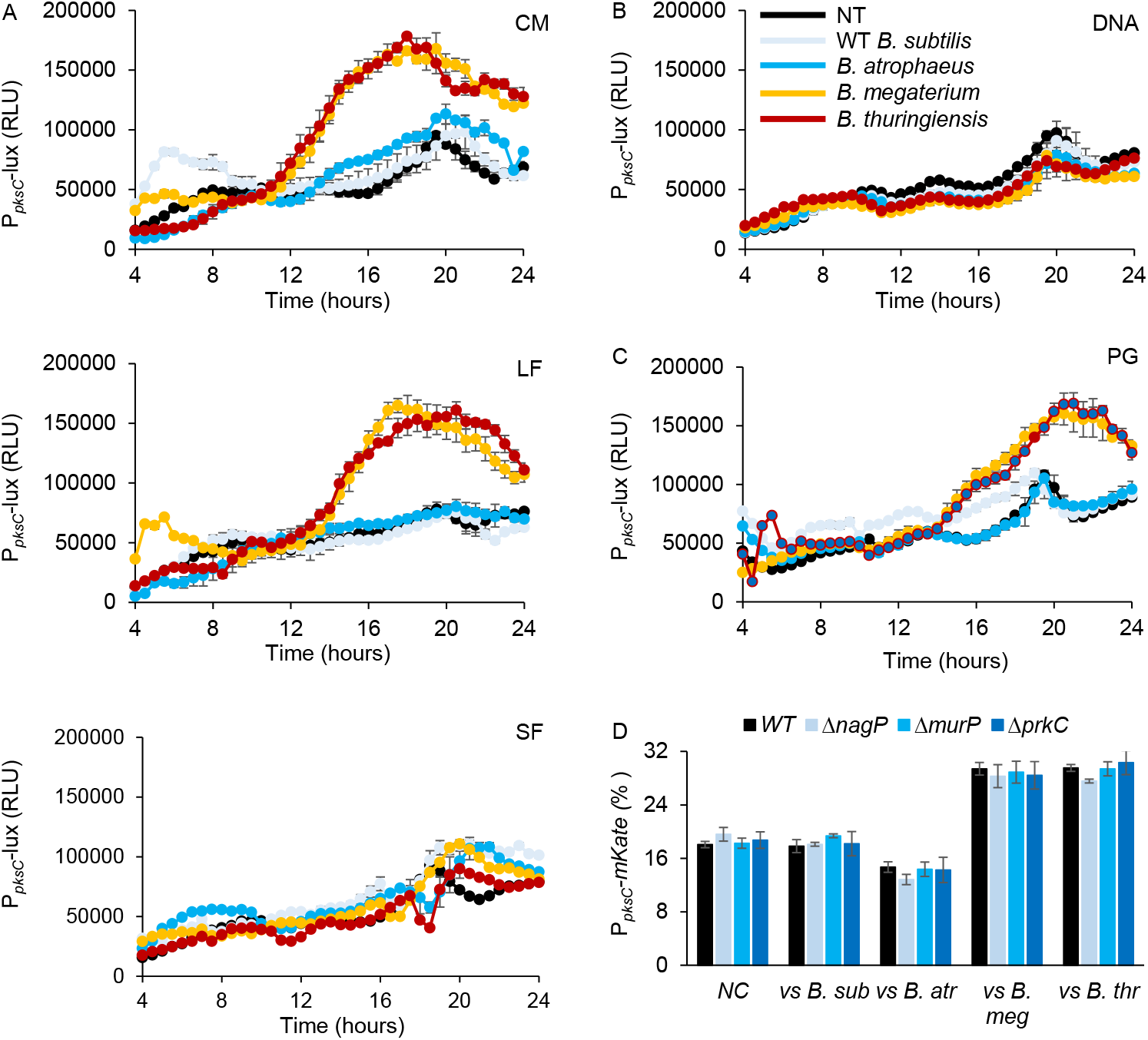
Distinguishing Self and Non-Self Regulates the Transcription from the Promoters of NRP Biosynthetic Clusters. **A)** Analysis of luciferase activity in *B. subtilis* cells harbouring P*_pksC_*-*lux* (bacillaene) reporter. Luminescence was monitored in B4 medium, supplemented either with DDW control or with 15% v/v of the conditioned medium (CM), or an equivalent amount of the conditioned medium fractionated to generate the LF (>3kDa) and SF (<3kDa) fractions. The donors of the conditioned media are indicated **B)** P*_pksC_*-*lux* (bacillaene) reporter activity in the presence of either purified gDNA or purified peptidoglycan (PG) of indicated strains. **C)** The parental reporter strain for P*_pksC_*-*mKate* (bacillaene) and its indicated mutants were analysed during either growth in isolation (NC) or at indicated interactions using flow cytometry. Data were collected from 24 h post inoculation. Y-axis represents fluorescent populations (100,000 cells were counted). Experiments were performed in triplicates, a representative example is shown from three independent experiments.

To identify the secreted factor which serves to activate the NRP transcription, we first separated the conditioned medium of self and non-self-competitors by size. A fraction >3Kda was sufficient to activate the transcription from both *srfAA* and *pks* promoters (Figure 5A and S8), while the smaller fraction was inert (Figure 5B). Therefore, the non-self-discrimination mechanism was by either a protein or a polymer. To exclude the involvement of secreted proteins and peptides, we treated the conditioned medium with proteinase K, a potent protease. As shown, the treatment did not reduce the potency of the conditioned medium (Figure S9). Similarly, purified exopolysaccharides (EPS) (Figure S10) failed to induce the transcription from *srfAA* and *pks* promoters. gDNA slightly induced transcription from *srfAA* promoter but the mild induction was comparable between self and non-self DNA (Figures 5B and S11). Thereof, a small molecule pheromone, EPS, proteins or extracellular DNA did not mediate non-self-recognition.

Peptidoglycan fragments have also been reported to function as signalling molecules that trigger adaptive responses. For instance, low concentration of muropeptides released by actively growing *B. subtilis* cells act as potent germinant of *B. subtilis* spores ^48^ Therefore, we tested whether non-self peptidoglycan fragments are sufficient to trigger non-self recognition of competing *Bacillus* species. Indeed, purified peptidoglycan from *B. megaterium* and *B. thuringiensis* but not *B. subtilis and B. atrophaeus* was sufficient to induce the expression of NRPs biosynthetic clusters at concentrations with little or no effect on cell growth (Figures 5B and S12).

We then explored which known regulators of NRPs antibiotics could act downstream of PG signals. In *B. subtilis*, canonical response to secreted PG is mediated by the PG sensors PrkC kinase^49^, recognizing muramopeptides, MurP^48^ for Muramic acid and NagP^48^ for N-Acetylglucosamine. However, deleting those genes had no effect on the basal levels or the induction of the transcription from NRPS promoters in the presence of non-self (Figures 5C, S13 and S14).

Surfactin and bacilysin are both regulated by the two-component regulatory system ComP/ComA involved in a major quorum response pathway that regulates the development of genetic competence^2^. We found that deletion of ComA was sufficient to eliminate non-self recognition of surfactin and bacillaene promoters. Specifically, for surfactin expression the deletion of ComA eliminated both basal expression and non-self recognition, and for Bacillaene the deletion eliminated specifically non-self recognition (Figure 6A, B and S15). In contrast, deletions of Spo0A and DegU, the master regulators of sporulation and biofilm formation^50^, did not specifically eliminate the response to non-self peptidoglycan. CodY regulated basal expression levels of from bacillaene biosynthesis promoter, but did not trigger non-self recognition (Figure 5A).

**Figure 6:**
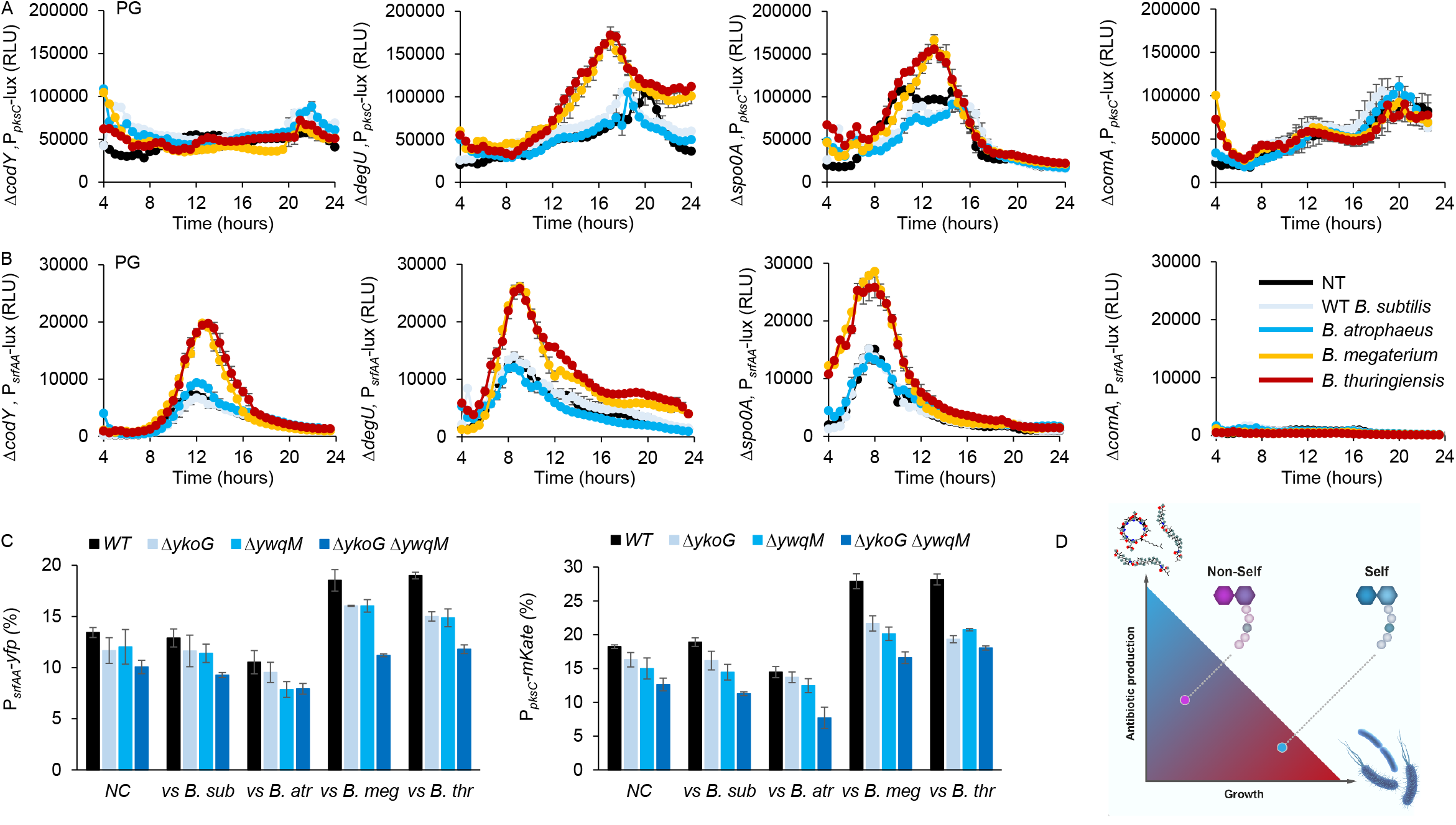
ComA and PG receptors mediate Non-Self Recognition. **(A-B)** Analysis of luciferase activity in *B. subtilis* parental strain (WT) and indicated mutants harboring P*_pksC_*-*lux* (bacillaene) (A) and P*_srfAA_*-*lux* (surfactin) (B) reporters. Luminescence was monitored in B4 medium (NT), and B4 medium supplemented with peptidoglycan (PG) from the indicated species. **C)** Indicated reporter strains for P*_pksC_*-*mKate* (bacillaene) and P*_srfAA_*-*yfp* (surfactin) and indicated mutants were analyzed for positively expressing populations either alone (NC) or at indicated interactions using flow cytometry. Data were collected from 24 h post inoculation. Y-axis represents fluorescent populations (100,000 cells were counted). Experiments were performed in triplicates. **D)** A conceptual model to the conflict between growth and NRPs production, and how it is resolved by the transcriptional regulation of antibiotic production.

*B. subtilis* genome has several homologues for known PG receptors^51^. YkoG, an homologue of the PG sensor BlrB ^48^ and YwqM, an homologue for the PG sensor AmpR ^48^ seem to be involved in NRPS transcriptional regulation. Although a deletion of a single homologue was insufficient to eliminate non-self PG recognition, a combination of both receptors was sufficient to significantly reduce non-self recognition in competing colonies (Figures 6C and Figure S16). Collectively, these results suggest that the ComA pathway is acting jointly with the PG sensory machinery (YwqM and YkoG) to fine-tune antibiotic production.

## Discussion

Microbial populations in natural environments secrete various secondary metabolites, such as antibiotics, digestive enzymes, quorum sensing inhibitors, low molecular weight compounds like hydrogen peroxide ^10, 20–23^, *etc*.

The primary model of our study is the plant biocontrol agent and probiotic bacterium *B. subtilis* ^2,52^. Various wild strains of *B. subtilis* and related species isolated from the soil are known to defend their host from diverse fungal and bacterial pathogens ^21,53^. These probiotic properties are largely mediated by the formation of non-ribosomal peptides ^54,55^.

In *B. subtilis*, a boundary is formed between kin and non-kin strains during swarming^11^, and this self-recognition was shown to involve contact dependent inhibition ^9^, while in *Proteus mirabilis* self recognition involved type VI secretion systems ^56,57^. It was previously hypothesized that direct cell damage (interference competition) activates bacteriocins and antibiotics^58^. Consistently, antimicrobials were also suggested to play a role in stabilizing communities composed of kin strains in *B. subtilis* ^9^. However, no specific antibiotic was sufficient for boundary formation between related and unrelated strains and the mechanism for non-self recognition remained to be determined.

Previous studies demonstrated that during microbial competition, numerous soil bacteria utilizes NRP antibiotics to ward off the invading microbes and thus protect their niche ^9,10,24,25^. Although molecular mechanisms of synthesis of the NRPs surfactin, bacillaene, bacilysin and pliplstatin is largely resolved ^59^, it remains unknown whether these antibiotics interact. Furthermore, bacteria that produce these antibiotics are subject to a fitness cost lowering their growth rate. Therefore, we used a genetically manipulatable antibiotic producer to study how different antibiotics interact, and to understand their regulation in the light of microbial competition and the conflict between antibiotic production and growth.

We found that the production of NRPs was specifically activated during competitions with sensitive species. Our results demonstrated that NRP antibiotics act as synergistic pairs (surfactin and bacillaene) or additively (with plipstatin and bacilysin) during interspecies competition. All antibiotics toxicity was specific towards competing species phylogenetically distant from the *Bacillus* genus, but sharing the same niche.

We therefore asked whether the expression of these biosynthetic clusters results in a deleterious effect on their producers’ growth in isolation. Indeed, production and increased production of NRPS came with a cost for the bacterial growth. This trade-off generates selective pressure on the regulation of NRP promoters (Figure 6D), as antibiotic production becomes deleterious when encountering closely related (and resistant) community members. Using transcriptional reporters, we showed that *B. subtilis* increased the transcription of NRP antibiotics only when sensing a specific secreted signal from sensitive species. This suggests that soil bacteria, such as *B. subtilis*, can discriminating “self” from “non-self”, and reduce the burden of producing antibiotics when encountering resistant clade members. Cell communication and metabolic exchange are essential part of interspecies competition and lead to the regulation of various molecular elements ^26,27^. Here we demonstrate how a PG cue for distinguishing of “self” from “non-self” can serve to minimize the cost for antibiotic production.

An essential feature of *B. subtilis* and Gram-positive bacteria in general is their cell wall. It counteracts the high intracellular osmotic pressure, determines the cell morphology, serves as diffusion barrier and serves as first layer of defence against environmental threats ^60,61^. Fragments of peptidoglycan, a mesh-like structure in the envelope, were shown to be sensed to induce the germination of spores ^49^, virulence genes of the human pathogen *P. aeruginosa* ^62^ and antibiotic resistance genes ^63^. Furthermore, the soil bacterium *Streptomyces coelicolor* induces the production of antimicrobials following perception of NAG from similar species ^30,45^. Here, we report on a new complementary role for PG recognition: distinguishing self and non-self phylum members competing for the same niche. The normal, unmodified glycan strands of bacterial peptidoglycan consist of alternating residues of β-1,4-linked N-acetylmuramic acid and N-acetylglucosamine. Glycan strands become differentially modified by in different species by enzymes responsible for the N-deacetylation, N-glycolylation and O-acetylation of the glycan strands^64^. The composition of the interpeptide bridge also differs between different species^64,65^, making PG fragments an appealing source of information regarding potential competitors’ identity,

Our observation that ComA acts downstream to PG sensing is especially intriguing. ComA is the transcription factor regulating genetic competence, a process in which *B. subtilis* takes up foreign DNA ^66^ It is not required for basal expression from bacillaene biosynthesis promoter (Figure 6A). However, it was required to regulate antibiotic production genes expression. These results indicate that genetic competence is co-regulated with a peak of antibiotics, in the presence of similar but not identical competitors. If so, these networks may indicate a co-evolution between the regulation of antibiotic production and horizontal gene transfer: competitors are sensed by PG receptors, lyzed by the induced antibiotics, and the DNA of these competitors is used to increase genetic diversity.

Disguising members of related species and unrelated species is of high importance, as members of the same clade are often resistant to their own self-produced antibiotics, and therefore the production of antibiotics under these conditions is more hazardous than beneficial for the producers. Overall, our findings indicate how multispecies communities of similar genotypes are favoured during competition, due to compatibility in antibiotic production and sensitivity. Coupling antibiotic production and the identification of “non-self” competitors provides a simple new principle that shapes bacterial communities, which may be further used for engineering microbiomes for ecological, medical and agricultural applications.

## Material and Methods

### Strains and media

All the strains used in this study are listed in supplementary Table S1.

Selective media for cloning purposes were prepared with LB broth or LB-agar using antibiotics at the following final concentrations: 10 μg/ml chloramphenicol (Amersco), 10 μg/ml, 10 μg/ml spectinomycin (Tivan biotech), 10 μg/ml tetracycline (Amresco), 1 μg/ml erythromycin (Amresco) + 25 μg/ml lincomycin (Sigma). Starter cultures of all strains was prepared using LB (Luria-Bertani) broth (Difco). B4 (0.4% yeast extract, (Difco), 0.5% D-glucose and 0.25% calcium acetate (Sigma Aldrich) was prepared as described previously in ^42^.

### Interaction assay

A single colony of each strain was isolated on a solid LB plate, inoculated into 2 ml of LB broth and grown to a mid-logarithmic at 37 °C with shaking. Mid-logarithmic cultures of *B. subtilis* and of each competitor were spotted on B4 plates 0.4 cm apart. The plates were then incubated at 30° C for 48 h and biofilms colonies were photographed using stereomicroscope (Zeiss). For CFU analysis, biofilms were harvested at 48 h and sonicated using BRANSON digital sonicator, at an amplitude of 10% and a pulse of 5 sec to separate cells without compromising their viability. 100 μl of sonicated cell suspension was transferred to Griener 90 well plate (Sigma) and a serial dilution ranging from 10^-1^ to 10^-7^ was performed using PBS (Phosphate-buffered saline). 20 μl of sample was transferred from the dilution series on a LB plate and incubated at 30 °C overnight and then counted.

### Community Experiments

The rhizosphere community (RC) present in the soil adhered to roots of organic *Solanum lycopersicum* with thickness at <1–2 mm was collected, suspended in 30 ml of PBS, and oscillated at 200 rpm at 30°C for 2 h. The roots were removed from the suspension and the PBS containing the RS was stored at −80 °C, for further use. The community experiments were performed using B4 medium. Briefly, 300ml of B4 medium was inoculated with 300 μL of the RS. To this, *B. subtilis* WT, Δ*srfAA*, Δ*pks* and Δ*4* samples were added in the final volume of ≈ 1000 CFU/ml. The set up was then incubated at 30°C with shaking for 48 h. For the CFU count, 100 μL of samples were taken at 0 h prior to incubation and at 24 h and 48 h after incubation and transferred to on a B4 plate with relevant antibiotics (see supplementary table S1), in order to filter and count only the relevant *B. subtilis* WT, Δ*srfAA*, Δ*pks* and Δ*4* population.

### Conditioned medium preparation

Cells were grown to a mid-logarithmic phase of growth (OD=0.6-0.8). Cells were diluted 1:100 in 300 ml of B4 medium and grown at 30°C for 24 h in shaker incubator (Brunswick^™^ Innova^®^ 42). Cells were removed by a centrifugation and the growth media was filtered by 0.22μm filter (Corning). CM was stored at 4 °C for further use.

### Growth measurements

Cells were grown from a single colony isolated over LB plates to a mid-logarithmic phase of growth. Cells were diluted 1:100 in 300 μl of B4 medium of each well of a 96-well microplate (Thermo Scientific). Cells were grown, with agitation, at 30°C for 24 h, in a microplate reader (Synergy 2; BioTek, Winooski, VT, USA), and the optical density at 600 nm was measured every 30 min. Cells were either grown in the presence or absence of CM (15 % V/V) as indicated in each corresponding figure legend.

### Flow Cytometry Analysis

Starter cultures were spotted on B4 plates 0.5 cm apart. The plates were then incubated at 30° C and biofilms colonies were harvested at 24 h and sonicated using BRANSON digital sonicator, model 250 at an amplitude of 10% and a pulse of 5 sec. Directly after sonication, samples were diluted in PBS and measured using an LSR-II cytometer (Becton Dickinson, San Jose, CA, USA). The GFP and YFP fluorescence was measured using laser excitation of 488 nm, coupled with 505 LP and 525/50 sequential filters. While mKate fluorescence was measured using laser excitation of 561 nm, coupled with 600 LP and 610/20 sequential filters. A total of 100,000 cells were counted for each sample and flow cytometry analyses was performed using FACS Diva (BD biosciences). Overlays of the flow cytometry results were obtained using FlowJo software.

### Luminescence analysis

Strains carrying the indicated luminescence reporters were grown in either B4 medium (NT) or B4 medium supplemented with the relevant indicated fractions of the indicated strains. A 96 – well plate with white opaque walls and clear tissue culture treated flat bottoms (Corning) was used for the measurements. Measurements were performed every 30 min at 30 °C, using a microplate reader (Synergy 2; BioTek, Winooski, VT, USA). Luciferase activity was calculated as RLU/OD.

### Statistical analysis

All studies were performed in triplicates at least three separate and independent times. All graphs were generated using Microsoft excel 2016. Data were expressed as average values ± standard deviations of the means. Data were analysed by one way ANOVA with Tukey post-hoc testing. Phylogenetic distances between *B. subtilis* and competitors were calculated using Integrated Microbial Genomes (IMG) database.

Additional methods are available in the supporting information: Strain construction,

## Author Contribution

HM, IKG, and JF designed experiments; HM performed the experiments; HM, IKG, and JF analysed the data; HM, IKG, and JF provided methodologies; IKG and HM wrote the manuscript.

## Acknowledgements

The Kolodkin-Gal lab is supported by the Israel Science Foundation grant number 119/16 and ISF-JSPS 184/20 and Israel Ministry of Science – Tashtiot (Infrastructures) −123402 in Life Sciences and Biomedical Sciences. IKG is supported by the Angel-Faivovich Fund for Ecological Research, by an internal grant from the Estate of Albert Engleman and by a research grant from the Benoziyo Endowment Fund for the Advancement of Science and a recipient of Rowland and Sylvia Career Development Chair. We thank Dr. Yaron Antebi (Dept. of Molecular Genetics) for his critical reading of our manuscript.

## Supporting Information

Supporting Figures (S1-S16)

Supporting Tables (S1-S4)

Supporting Methods

Supporting References

## Supplementary Figures

**Figure S1:**
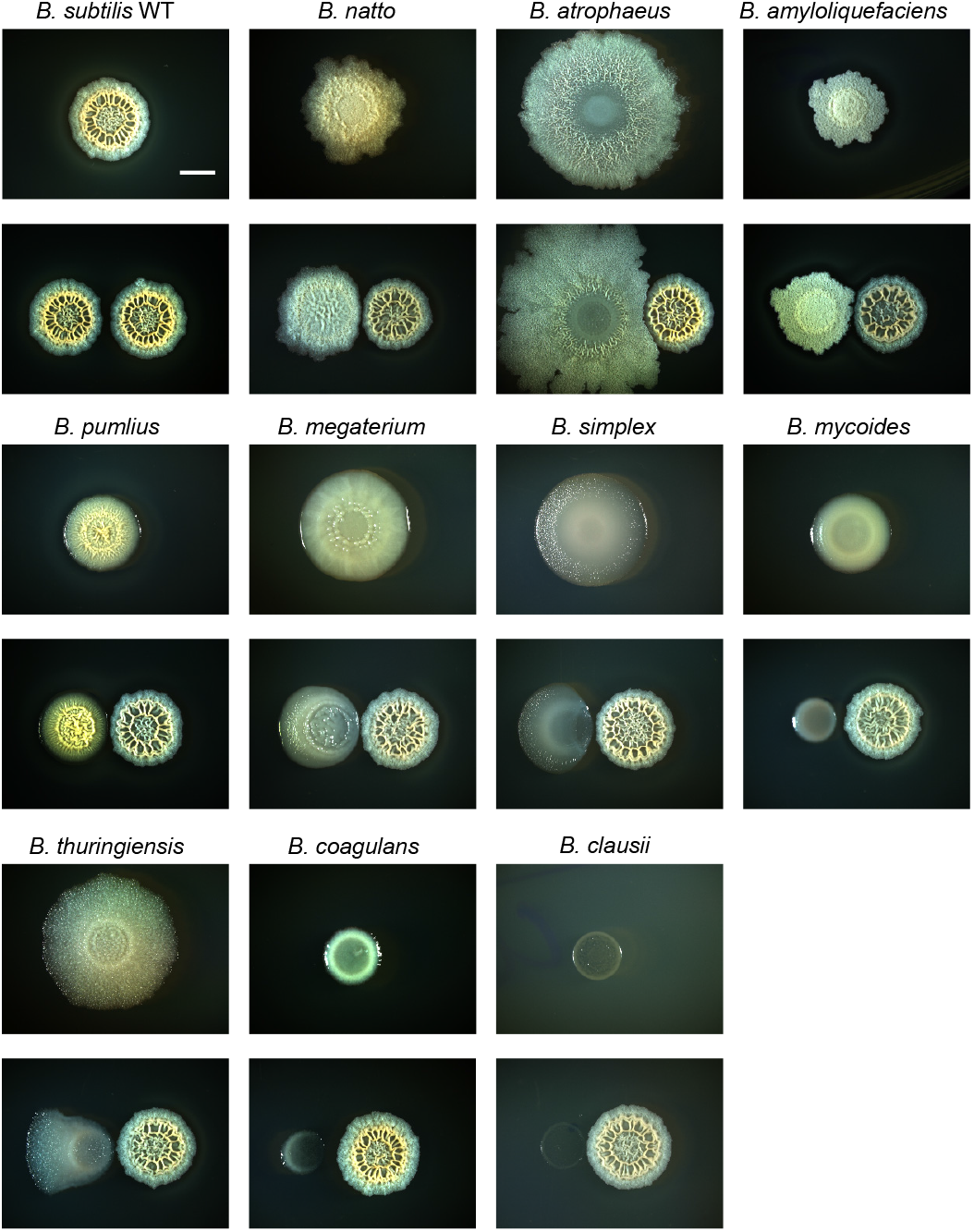
Growth in isolation and during competition of indicated *bacillus* species. For each species, shown is growth in isolation versus growth in interspecies competition against *B. subtilis* colony (below). Colony biofilms were inoculated at 0.4 cm apart and grown on B4 medium at 30° C. Biofilms colonies were imaged 48 hours post inoculation. Scale bar = 1mm.

**Figure S2:**
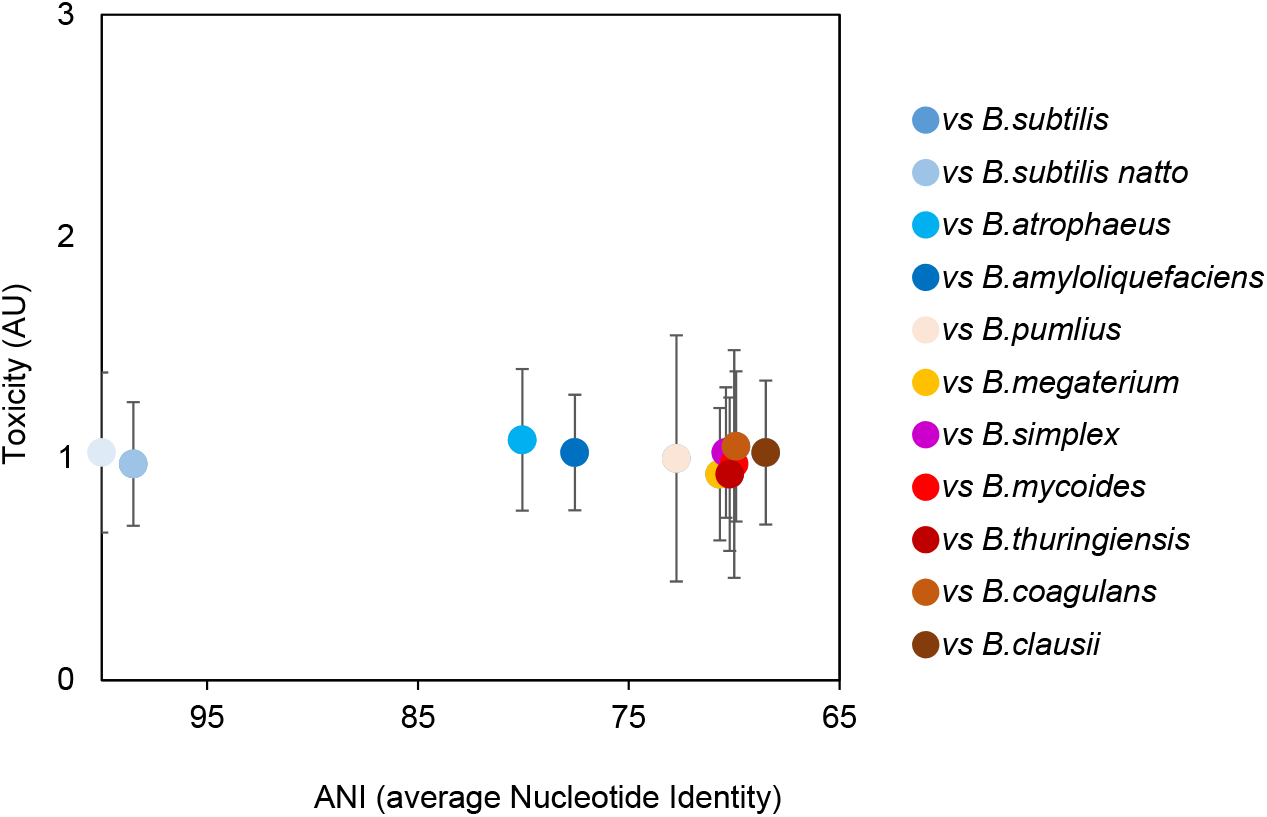
Toxicity of bacilli towards *B. subtilis* was evaluated. AU for toxicity were determined as the ratio of *B. subtilis* growth alone/growth in competition against bacilli: Biofilm cells were harvested prior to direct contact and colony forming units (CFU) were calculated alone, and during co-inoculation. X-axis represents the average nucleotide identity of competing *bacilli*.

**Figure S3:**
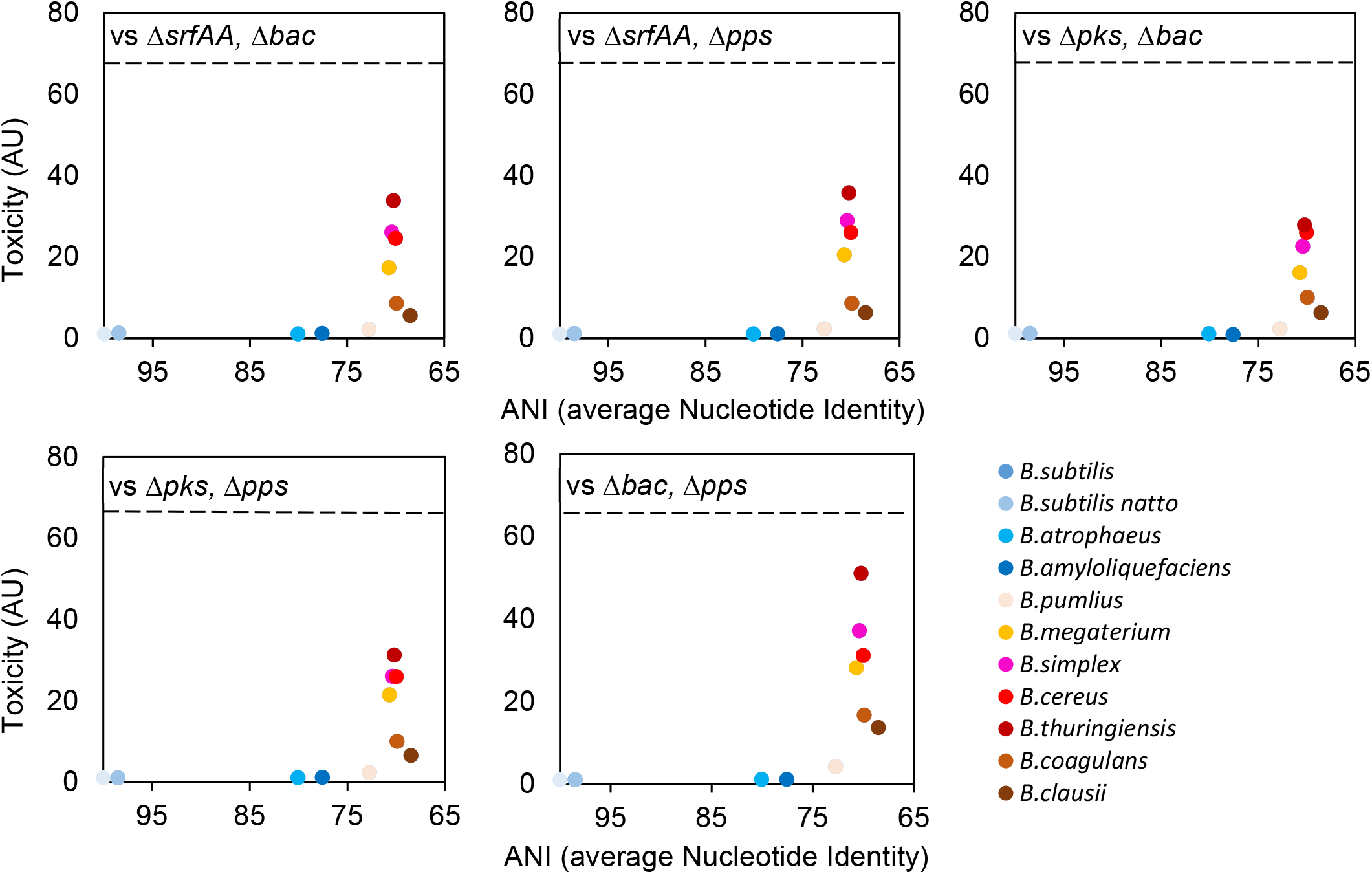
Toxicity of indicated mutants towards competitors was evaluated. AU for toxicity were determined as the ration of growth alone/growth in competition against *B. subtilis:* Biofilm cells were harvested prior to direct contact and colony forming units (CFU) were calculated alone, and during co-inoculation. X-axis represents the average nucleotide identity of competing *bacilli*. Dashed Line: The toxicity of the WT *B. subtilis*

**Figure S4:**
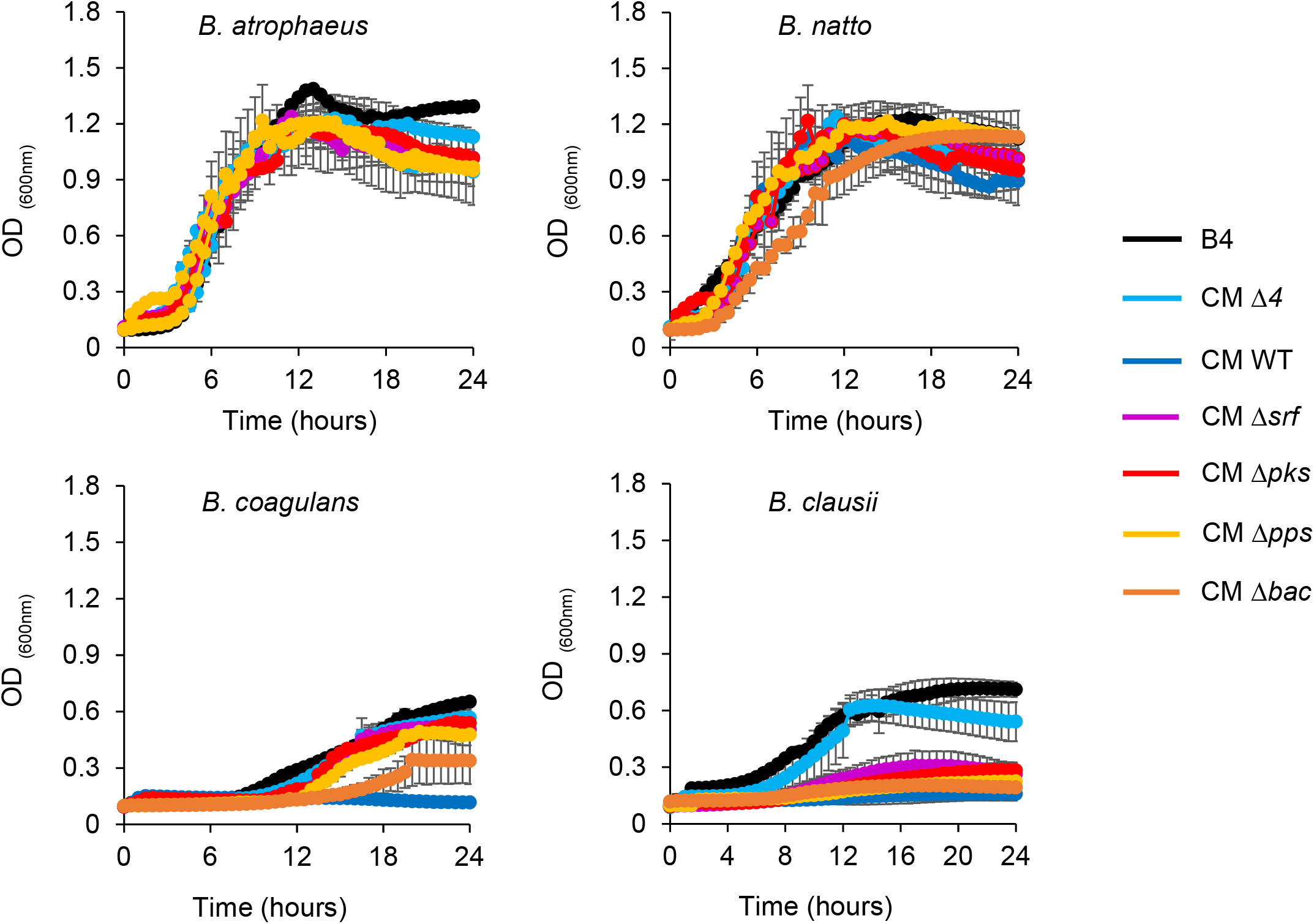
Planktonic growth of the indicated species was monitored either in B4 medium or B4 medium supplemented with CM of the parental strain (WT) 15% v/v or it’s indicated NRPs biosynthetic clusters mutants15% v/v.

**Figure S5:**
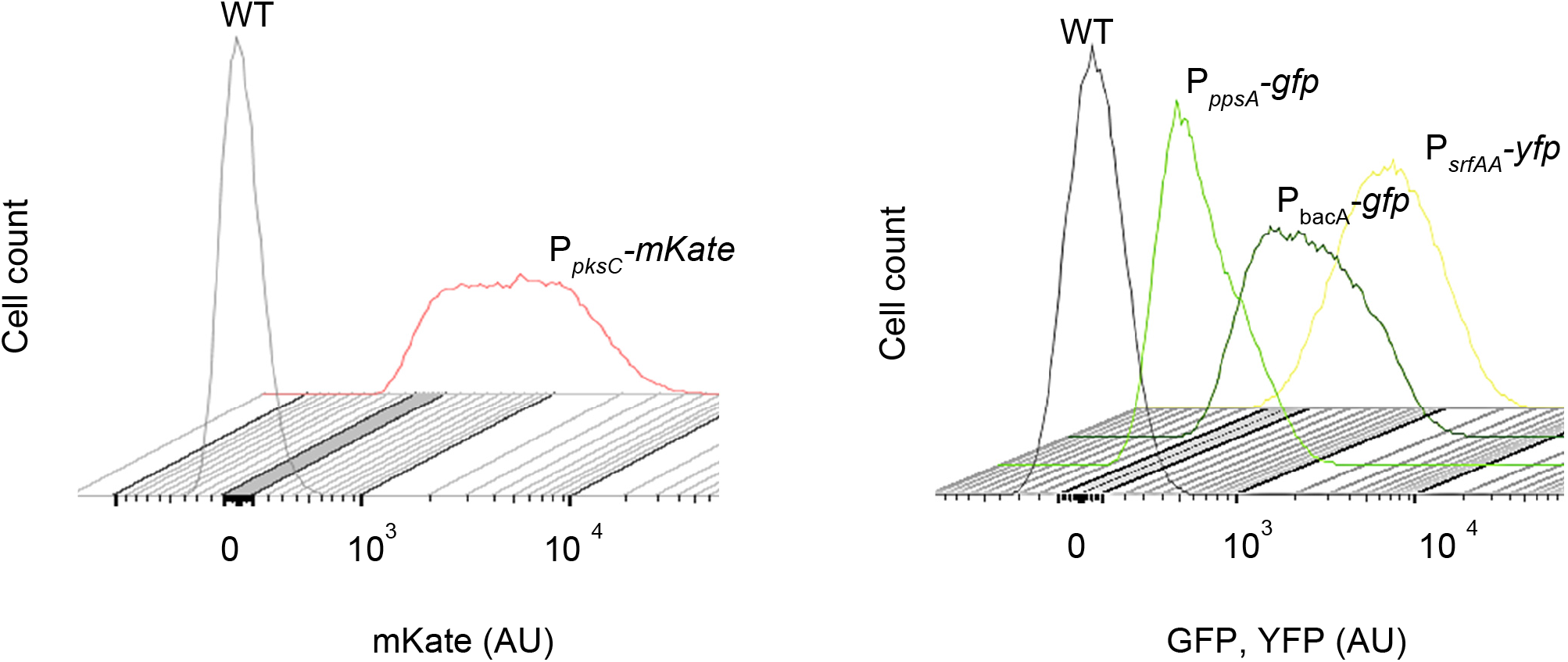
Flow cytometry analysis of a biofilm colony for indicated reporter genes. Colonies were grown on B4 medium and incubated at 30°C; flow cytometry analysis was performed at 24 h time point. Grey peak represents the negative control population of *B. subtilis* (no fluorescent reporter). The Y axis represents cell count (100,000 cells were counted), and X-axis is arbitrary units (AU) of fluorescence in log scale. Overlays comparing basal expression of P*_srfAA_*-*yfp* (surfactin), P*_pksC_ mKate* (bacillaene), P*_bacA_*-*gfp* (bacilysin) and P*_ppsA_* - *gfp* (plipastatin).

**Figure S6:**
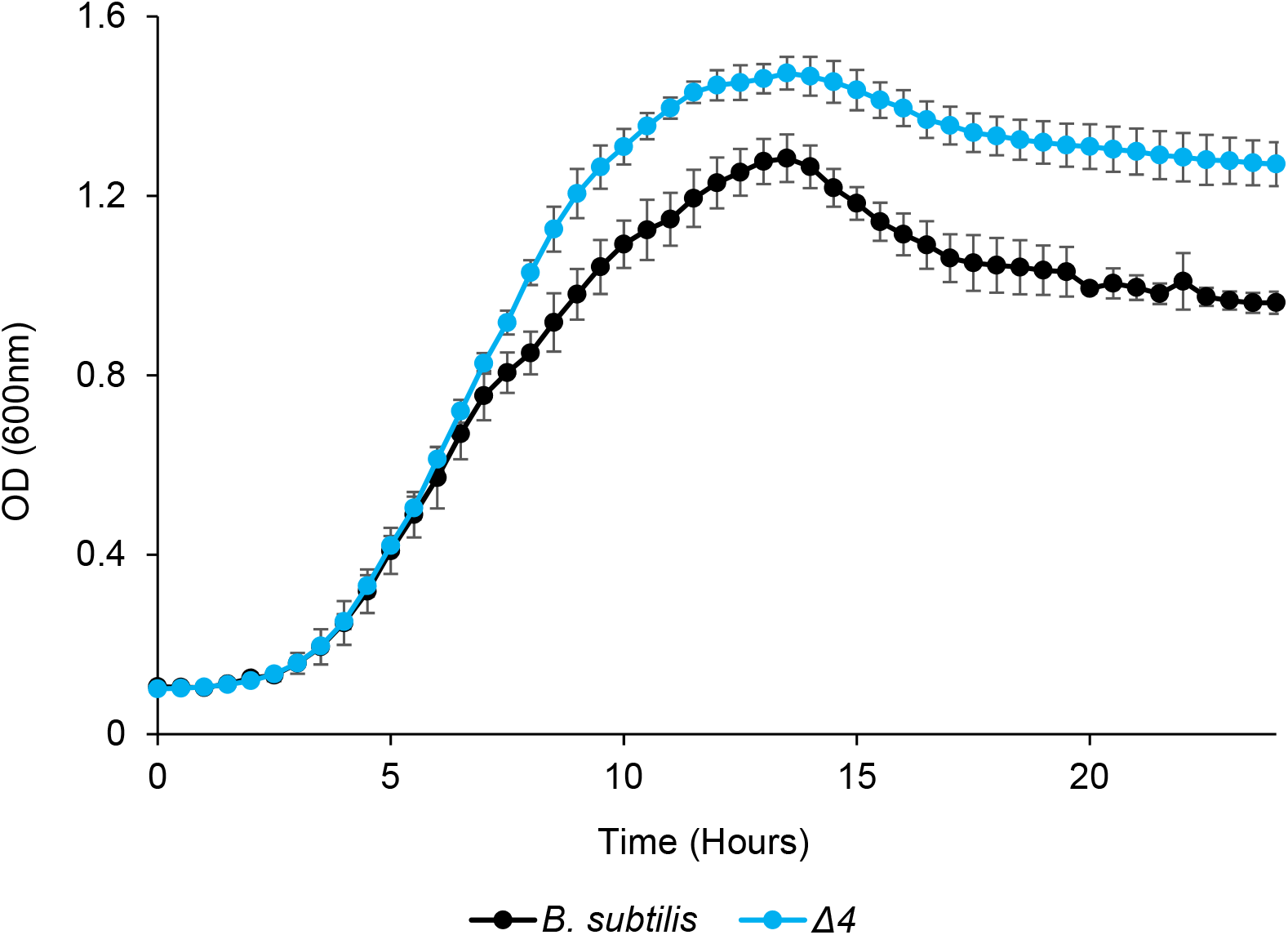
Planktonic growth of the B. subtilis WT and a quadruple mutant for all NRPS operons *srfAA- srfAD, pksC- pksR, bacA-ywfG and ppsA –ppsE* (Δ*4*).

**Figure S7:**
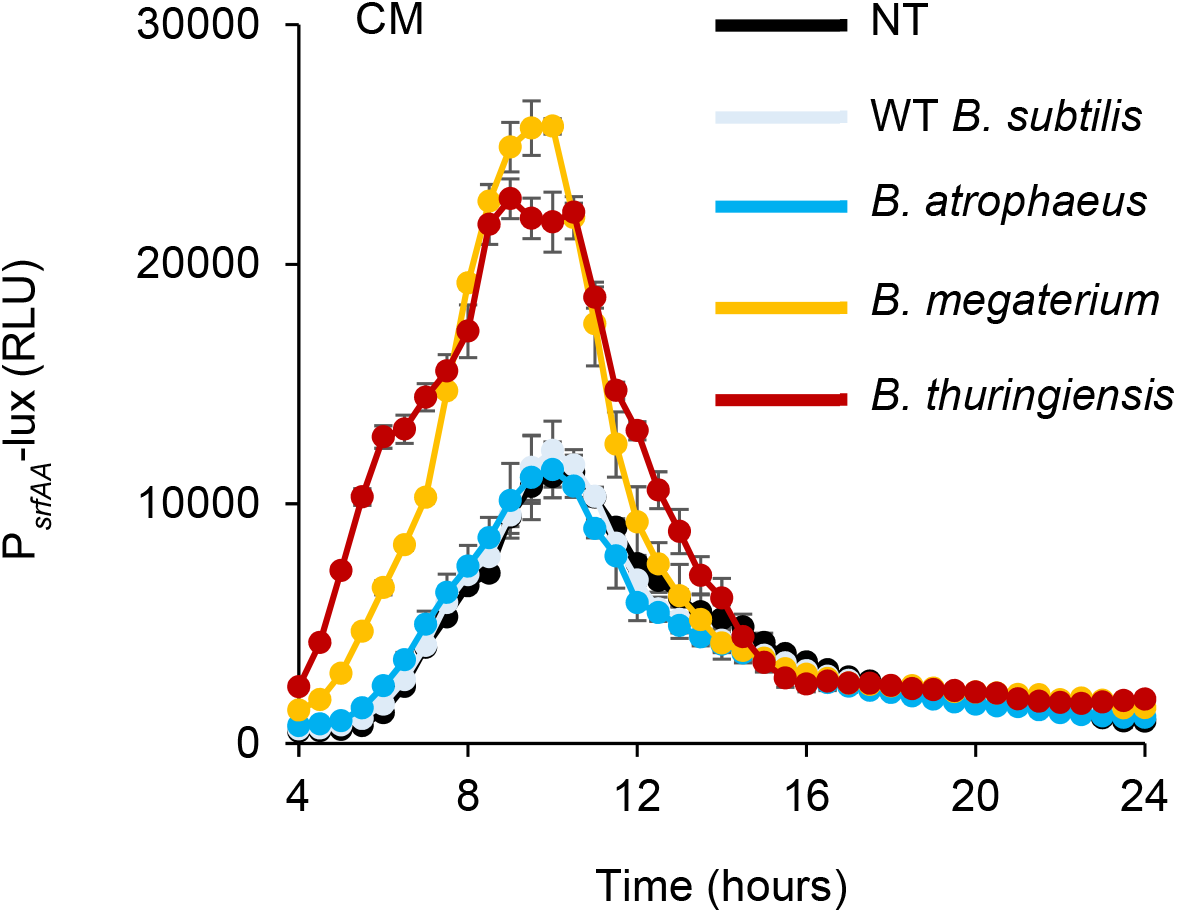
Analysis of the luciferase activity in a *B. subtilis* strain harboring P*_srfAA_*-*lux* (surfactin) reporter. Luminescence was monitored in B4 medium (NT), and B4 medium supplemented with 15% v/v of the conditioned medium (CM) from the indicated species.

**Figure S8:**
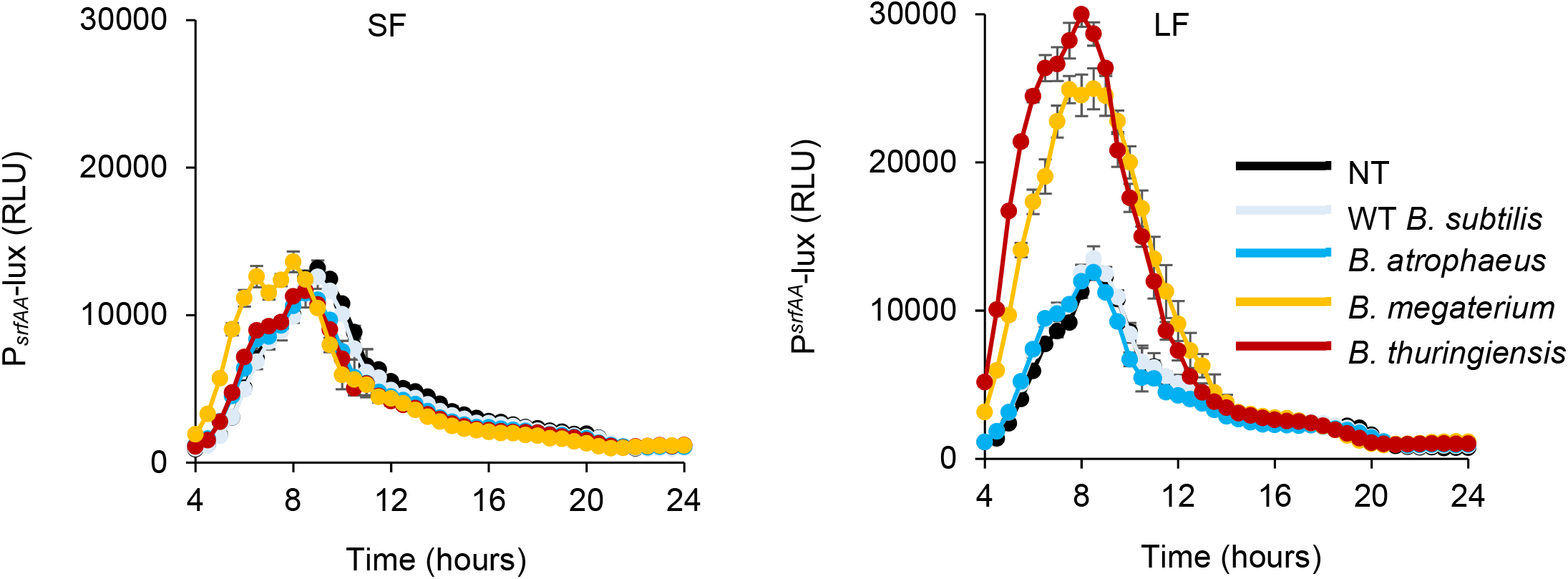
Analysis of the luciferase activity in a *B. subtilis* strain harbouring P*_pksC_*-*lux* (bacillaene) reporter. Luminescence was monitored in either B4 medium (NT), or B4 medium supplemented with equivalent amount of the dialyzed conditioned medium. The dialysis generated SF (<3kDa) and LF (>3kDa).

**Figure S9:**
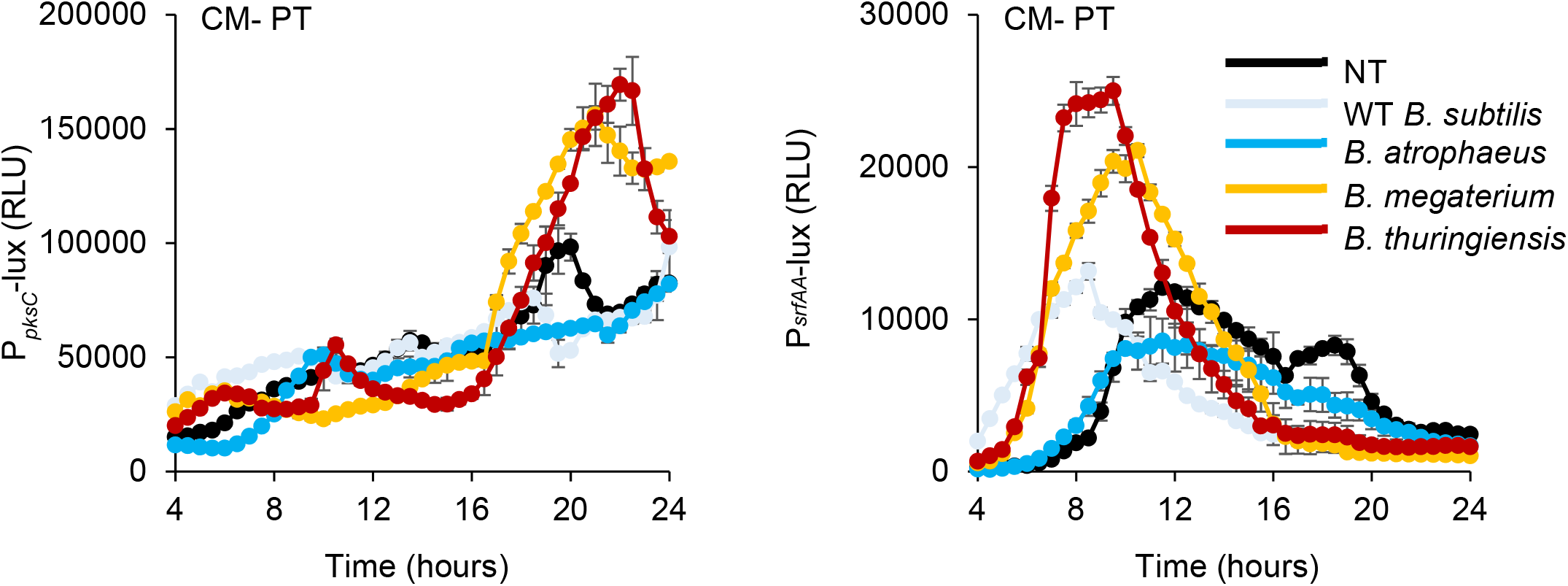
Analysis of the luciferase activity in a *B. subtilis* strain harboring P*_pksC_*-*lux* (bacillaene) and P*_srAA_*- *lux* (surfactin) reporter. Luminescence was monitored in either B4 medium (NT), or B4 medium supplemented with 15% v/v of protease treated (PT) conditioned medium (CM) from the indicated species.

**Figure S10:**
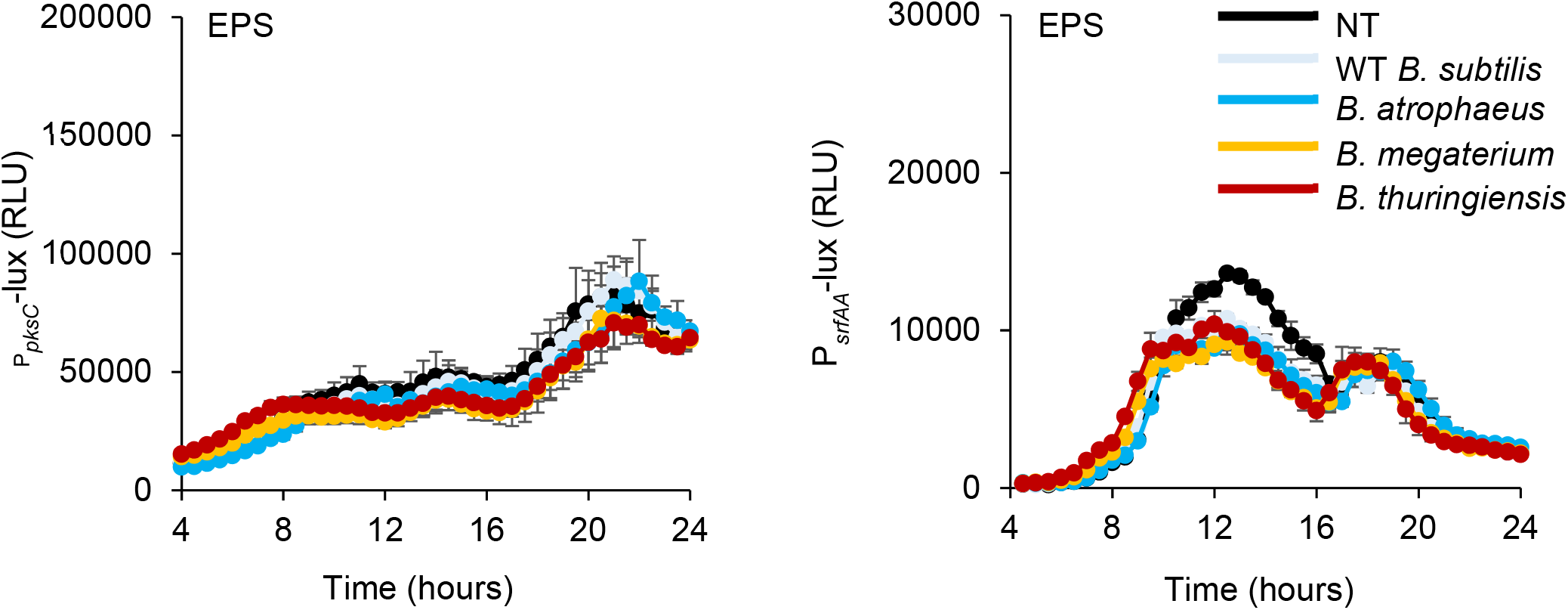
Analysis of luciferase activity in *B. subtilis* cells harboring P*_pksC_*-*lux* (bacillaene) and P*_SfAA_*-*lux* (surfactin) reporter. Luminescence was monitored in either B4 medium (NT), or B4 medium supplemented with purified EPS from the indicated species.

**Figure S11:**
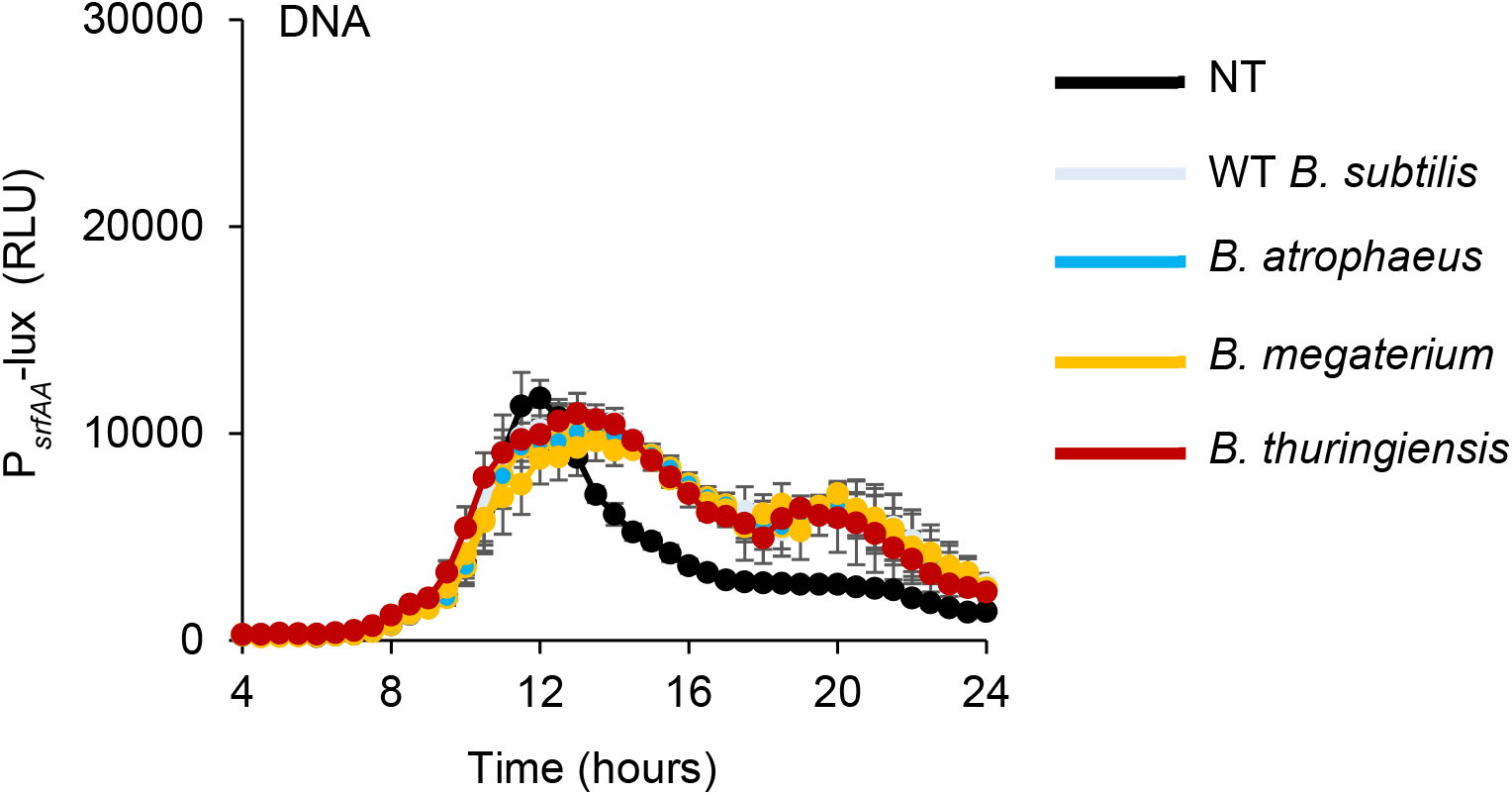
Analysis of luciferase activity in *B. subtilis* cells harboring P*_SrfAA_*-*lux* (surfactin) reporter. Luminescence was monitored in either B4 medium (NT), or B4 medium supplemented with gDNA from the indicated species.

**Figure S12:**
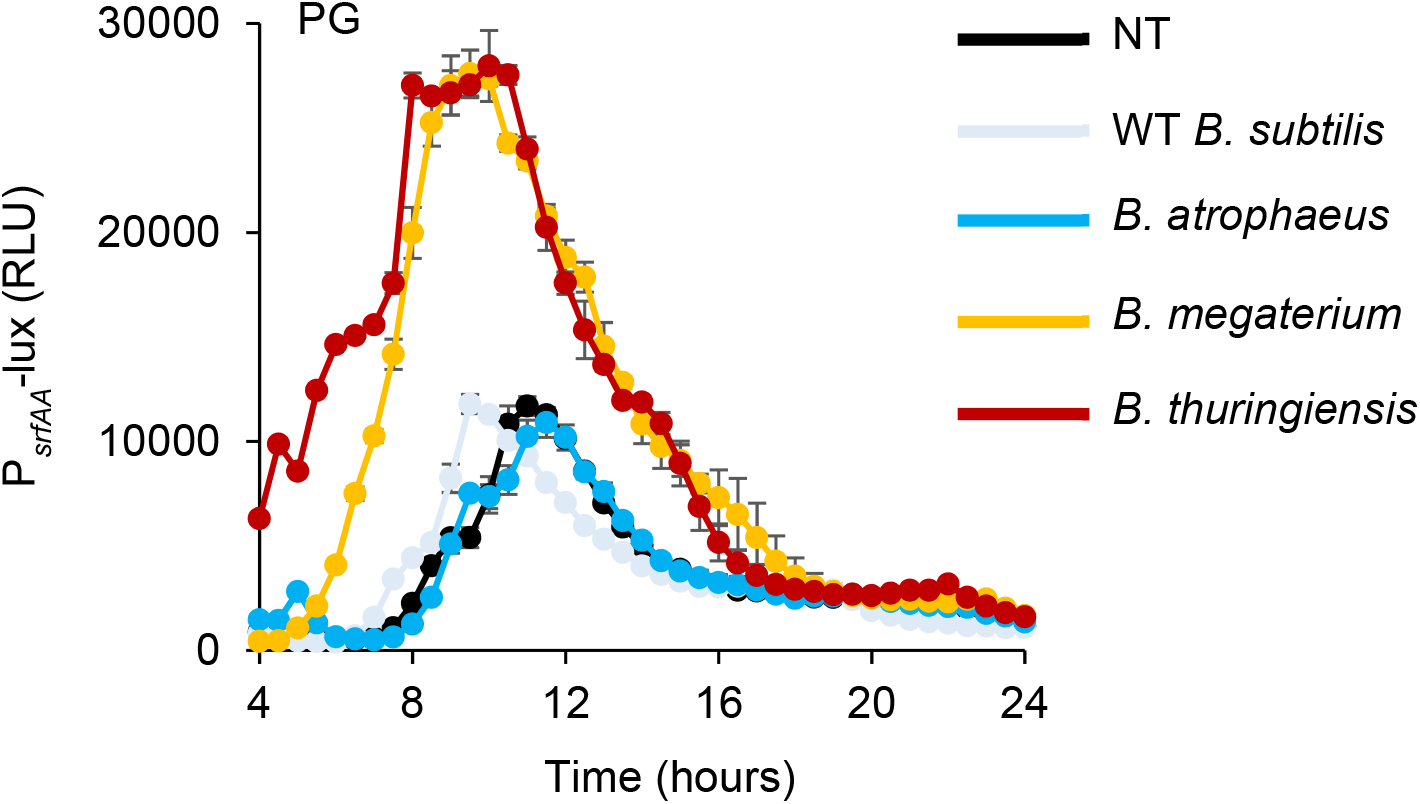
Analysis of luciferase activity in *B. subtilis* cells harboring P*_SrfAA_*-*lux* (surfactin) reporter. Luminescence was monitored in either B4 medium (NT), or B4 medium supplemented with purified peptidoglycan (PG) from the indicated species.

**Figure S13:**
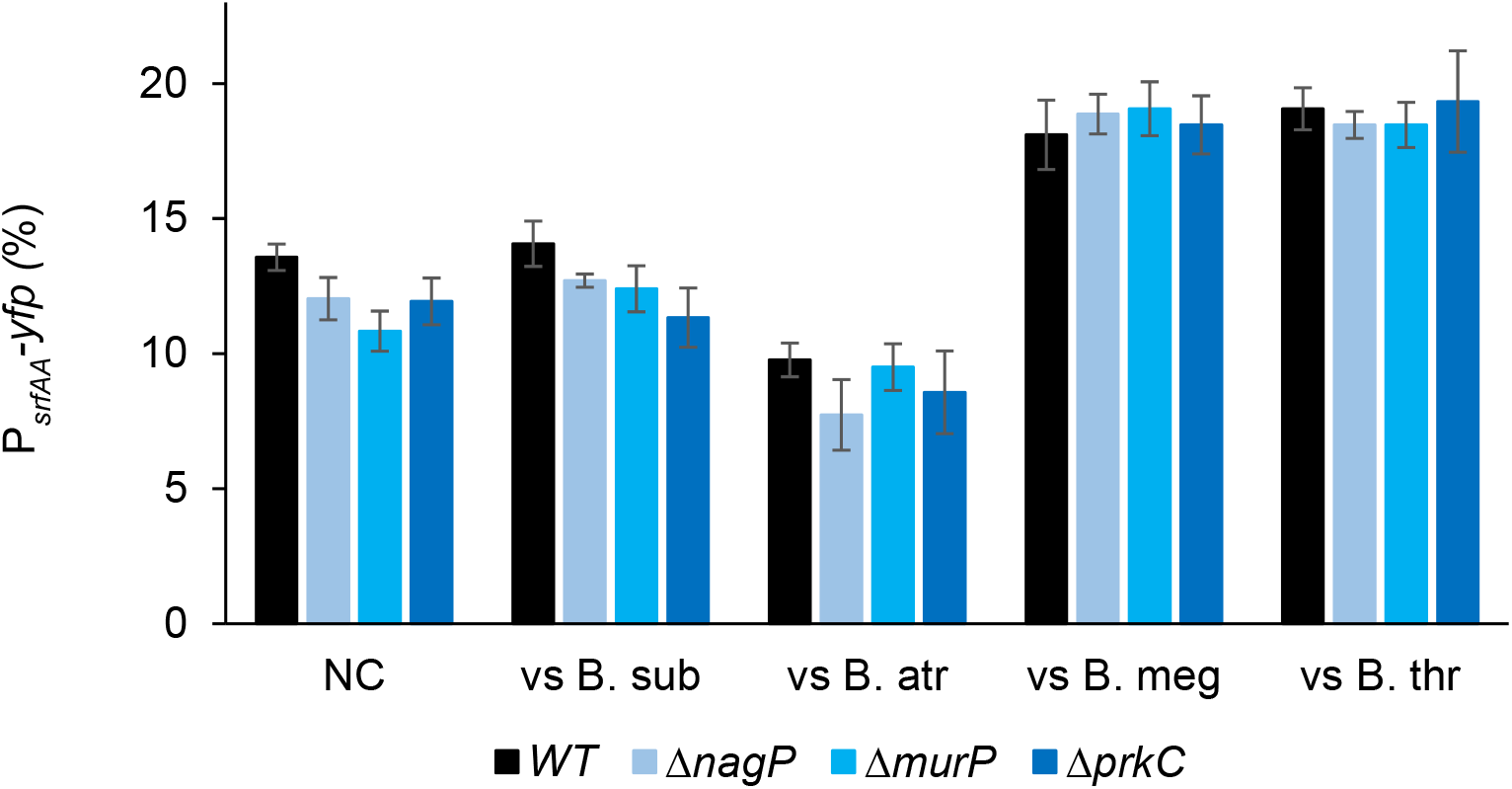
A parental strain P*_srfAA_*-*yfp* (surfactin) WT and its indicated mutants were analyzed either alone (NC) or in competition against with bacilli using flow cytometry. Colonies were grown on B4 medium and incubated at 30°C. Data were collected from 24 h post inoculation and prior to direct contact Y-axis represents fluorescent populations (100,000 cells were counted). Shown is the % of cells expressing the reporter.

**Figure S14:**
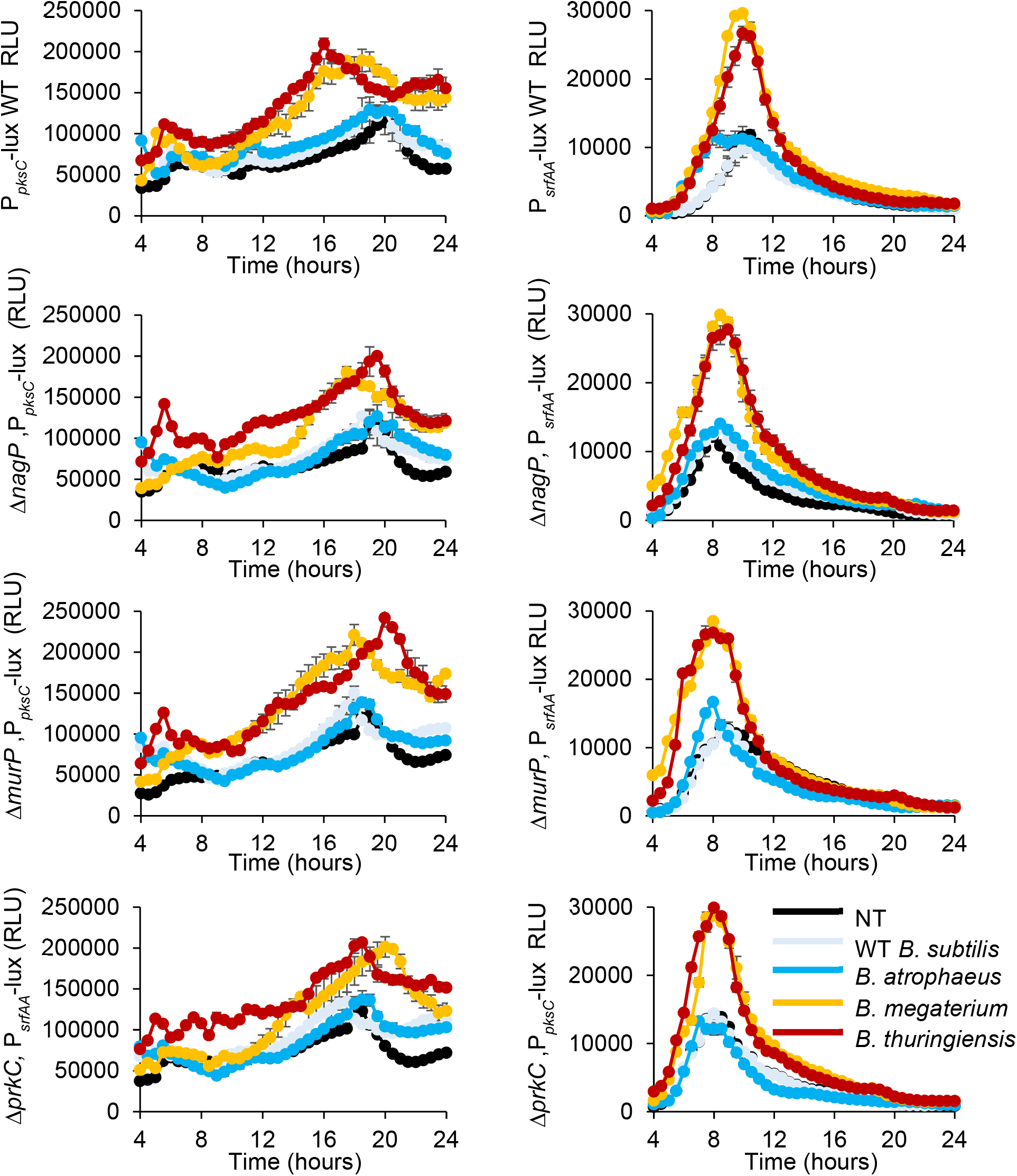
Analysis of luciferase activity in *B. subtilis* parental strain (WT) and its indicated mutants harboring P*_pksC_*-*lux* (bacillaene) and P*_srfAA_*-*lux* (surfactin) reporters. Luminescence was monitored either B4 medium (NT), and B4 medium supplemented with peptidoglycan (PG) from the indicated species.

**Figure S15:**
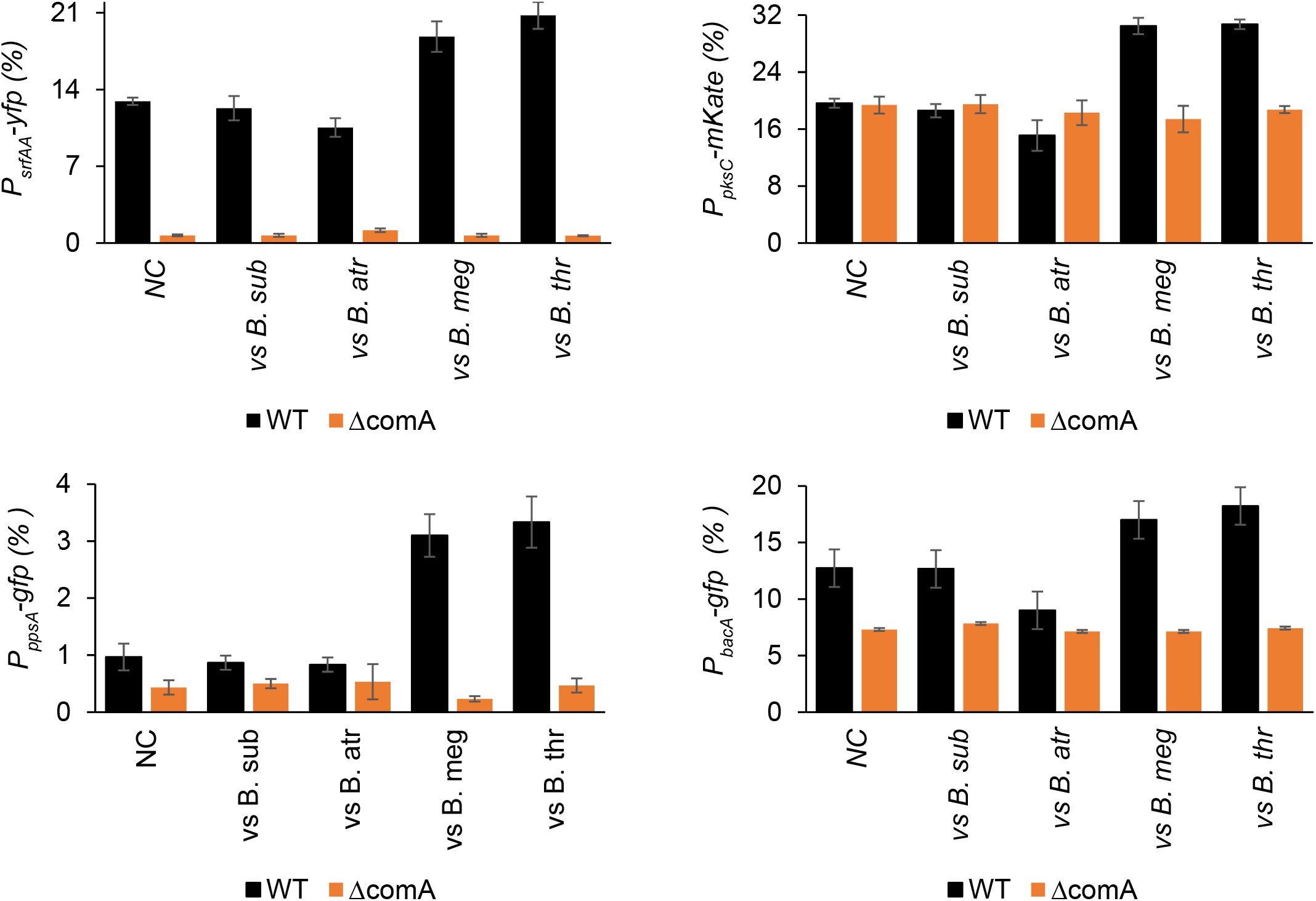
A parental strain (WT) carrying either P*_srfAA_*-*yfp* (surfactin), P*_pksC_*-*mKate* (bacillaene), P*_ppsA_*-*gfp* (plipastatin) or P*_bacA_*-*gfp* (bacilysin), and its Δ*comA* mutant were analyzed either alone (NC) or in competition against with bacilli using flow cytometry. Colonies were grown on B4 medium and incubated at 30°C. Data were collected from 24 h post inoculation and prior to direct contact Y-axis represents fluorescent populations (100,000 cells were counted). Shown is the % of positive cells.

**Figure S16:**
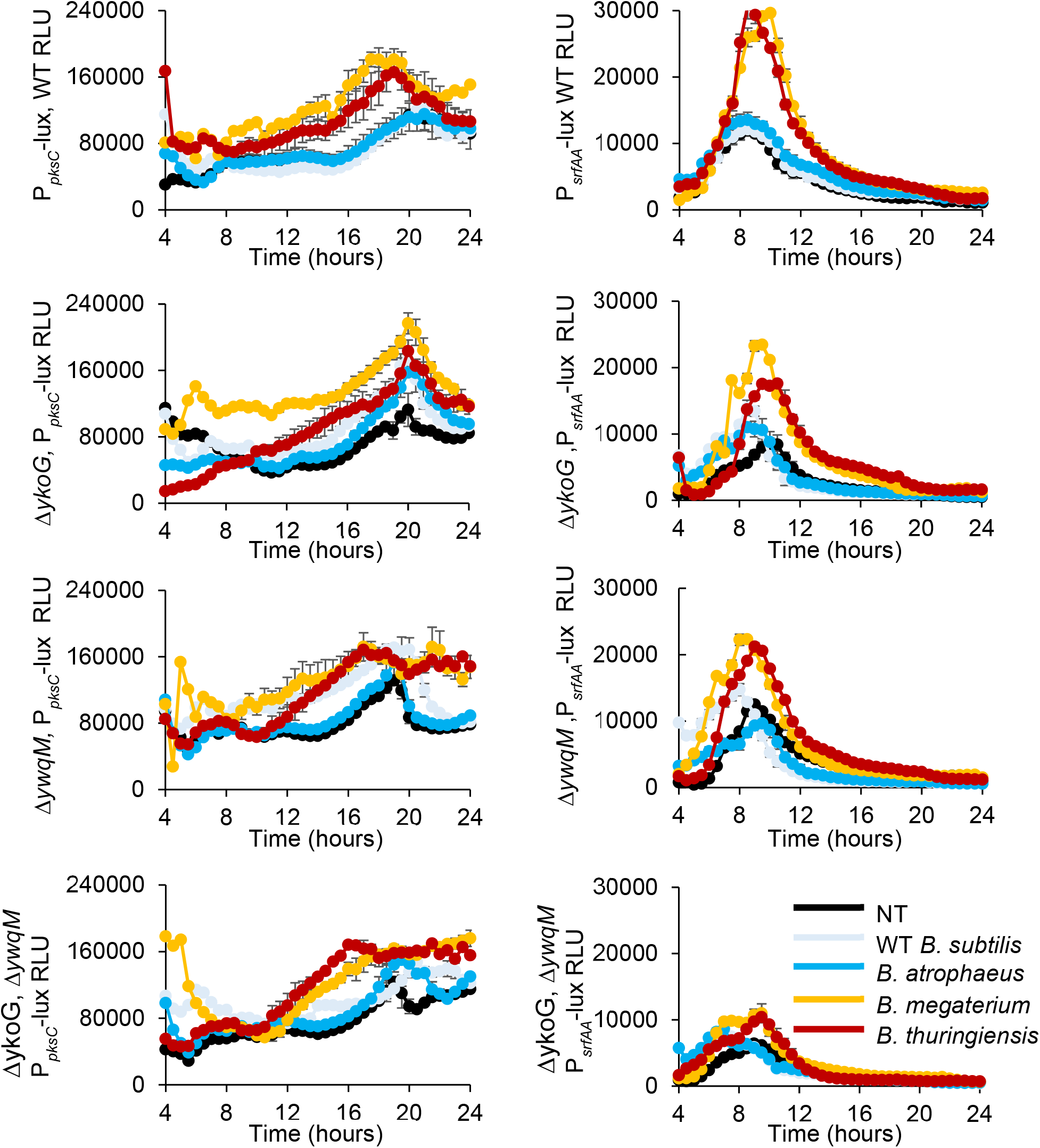
Analysis of luciferase activity in *B. subtilis* parental strain (WT) and its indicated mutants harboring P*_pksC_*-*lux* (bacillaene) and P*_srfAA_*-*lux* (surfactin) reporters. Luminescence was monitored either B4 medium (NT), or B4 medium supplemented with peptidoglycan (PG) from the indicated species.

**Table S1.**
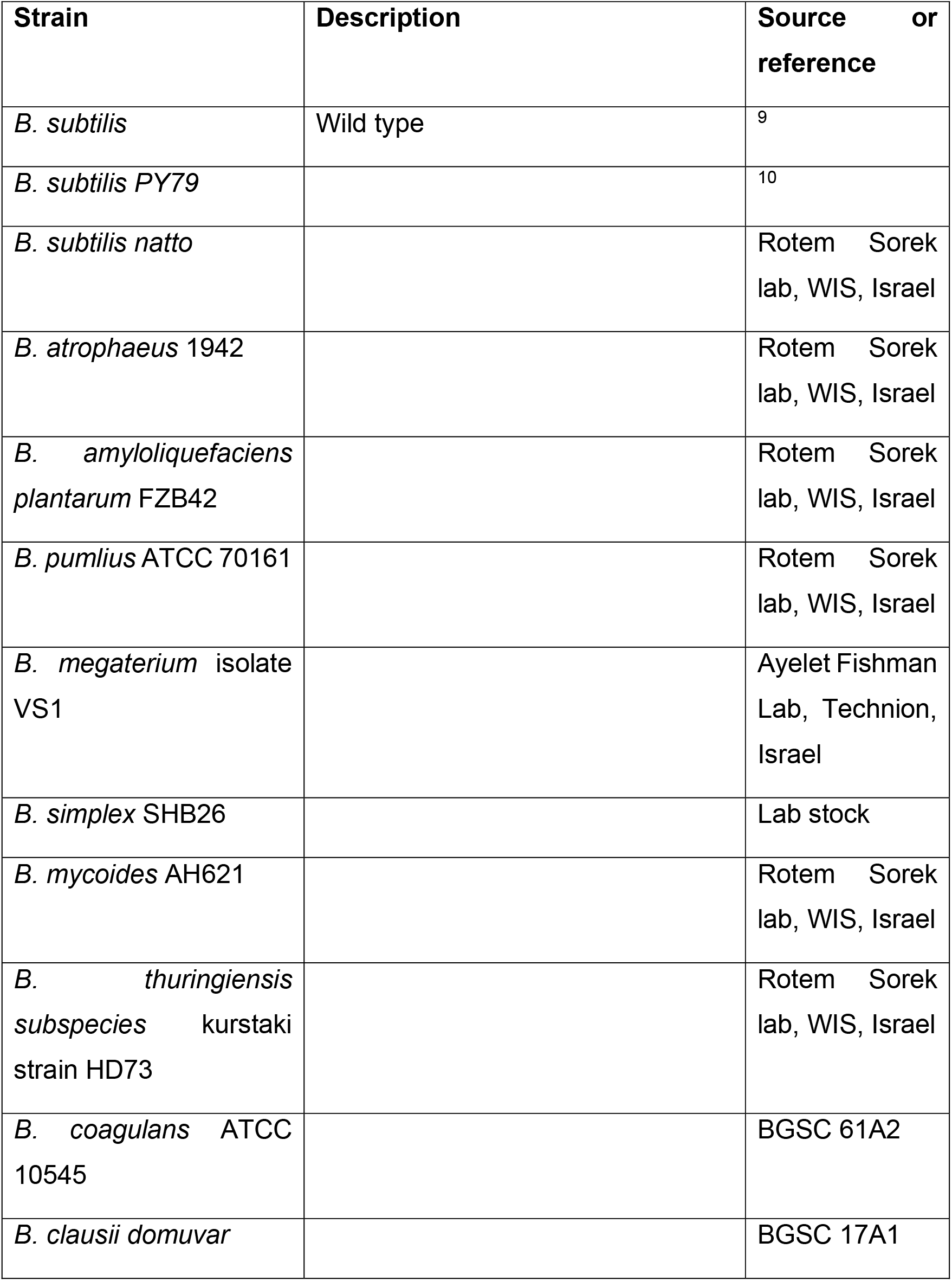
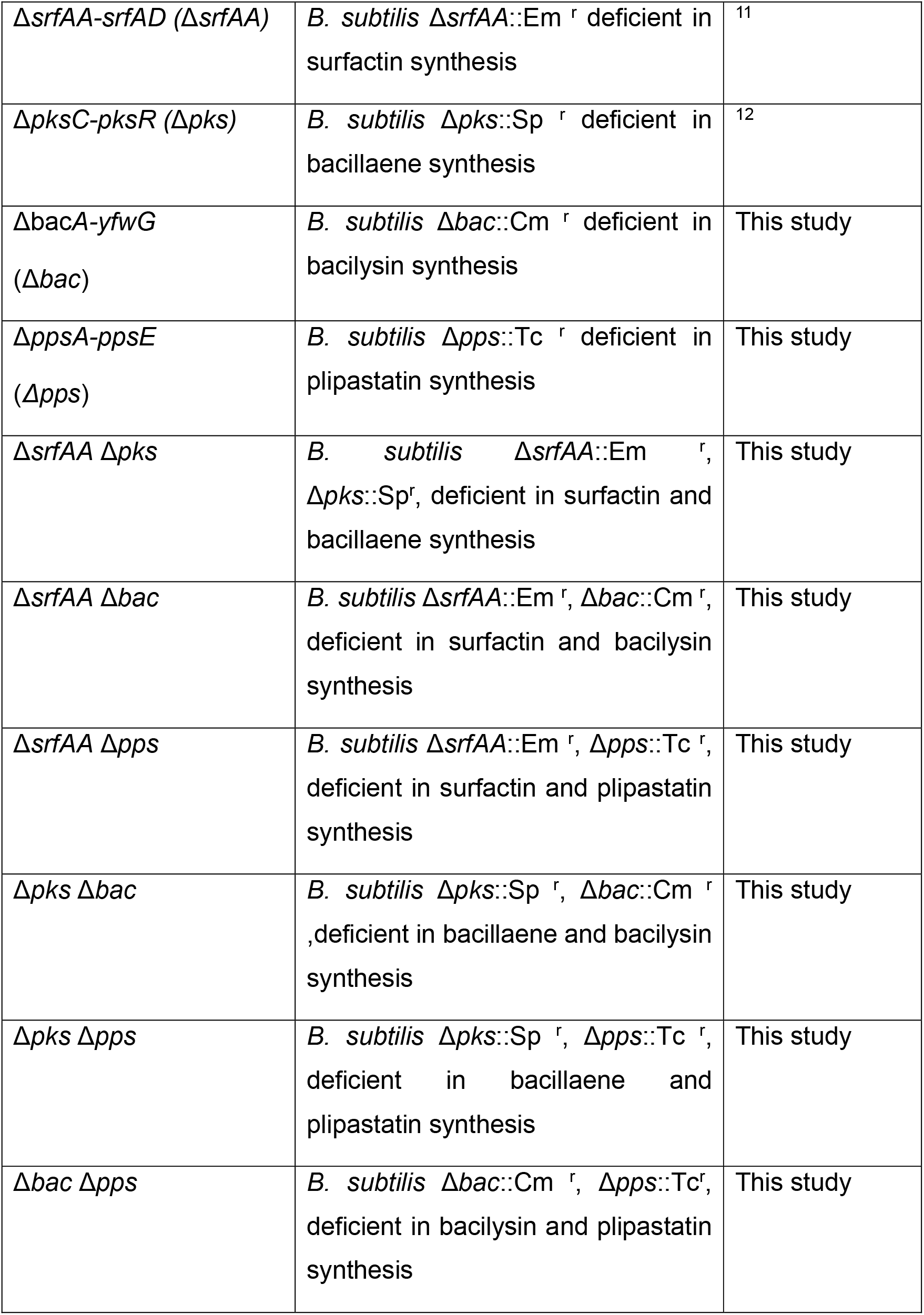
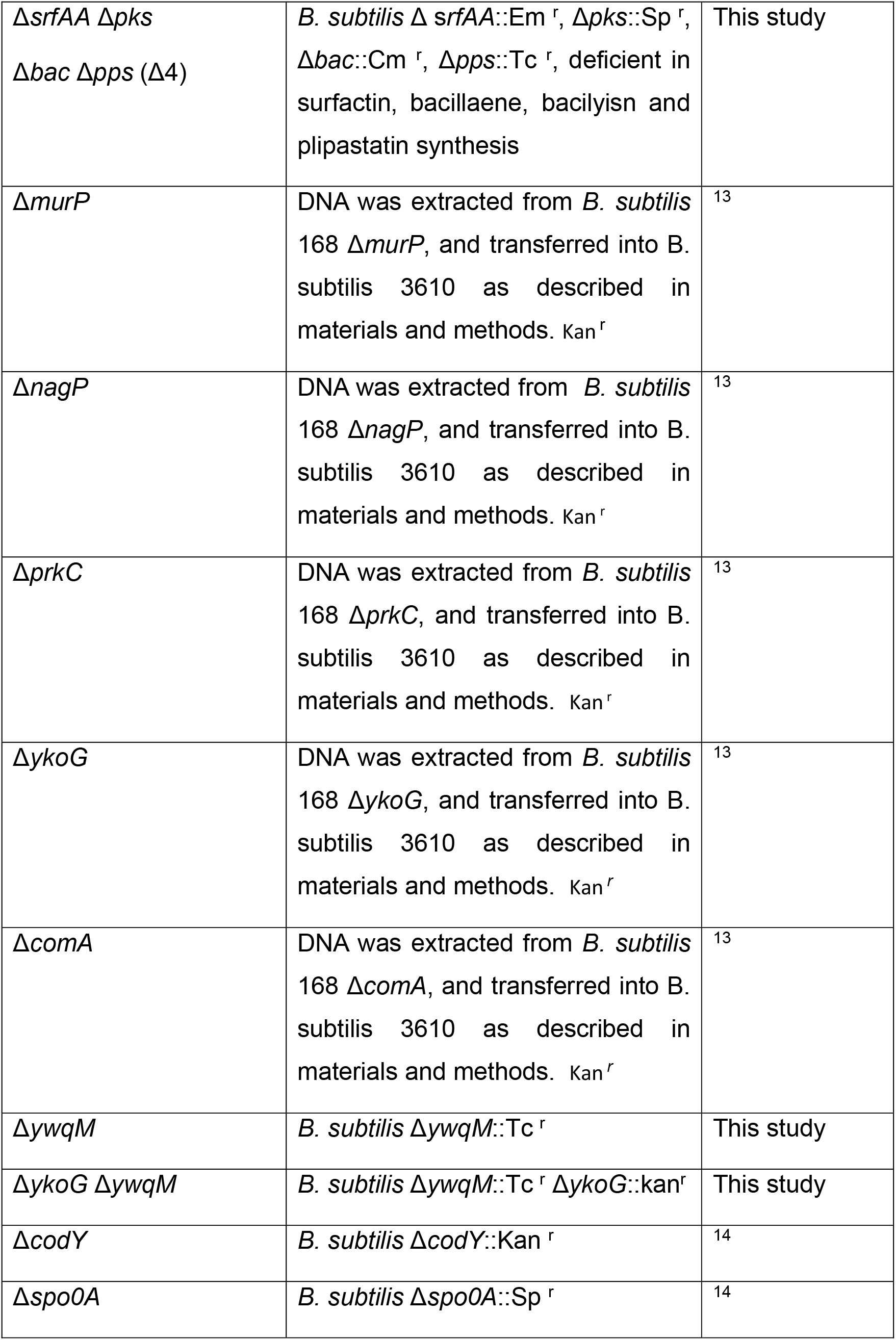
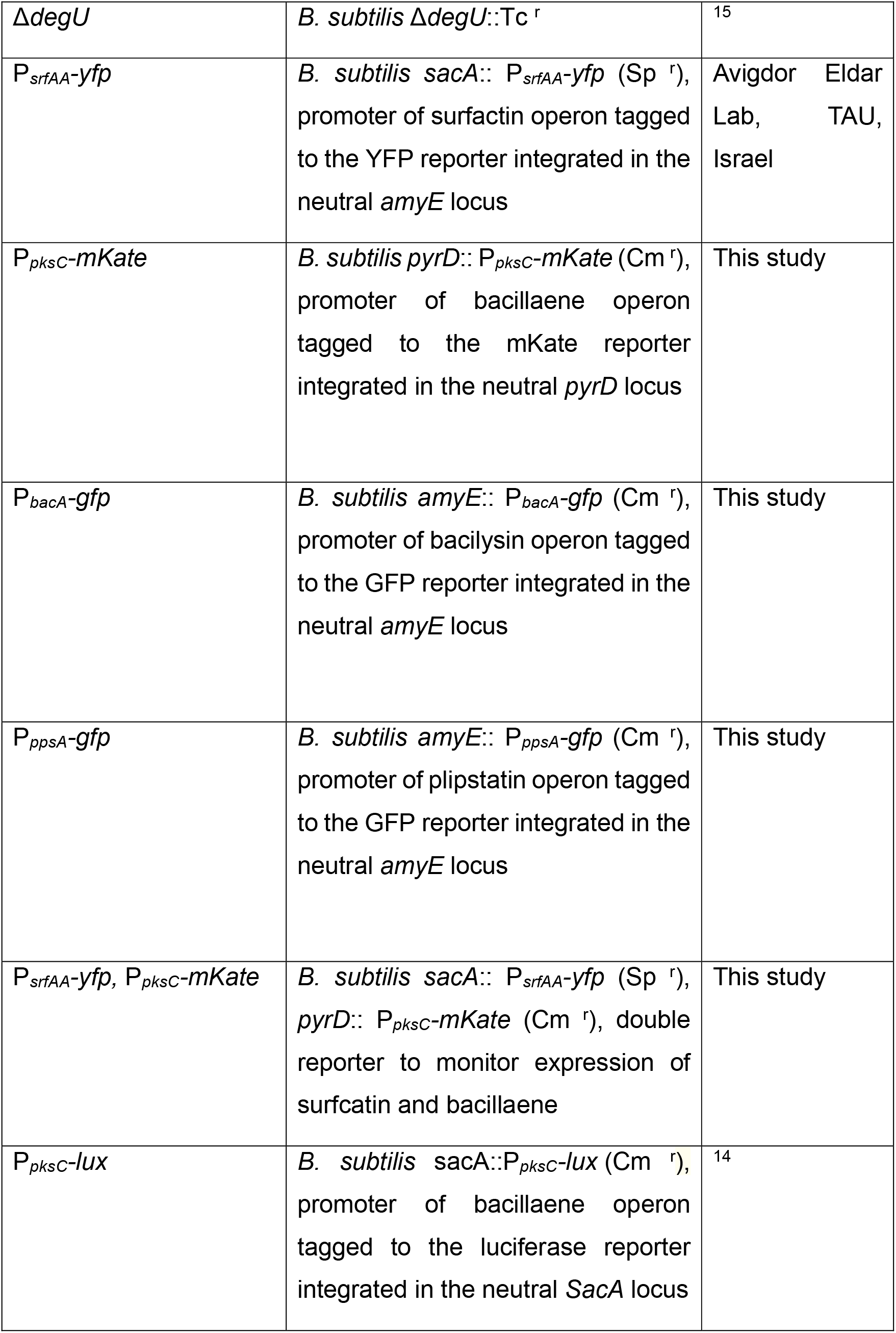
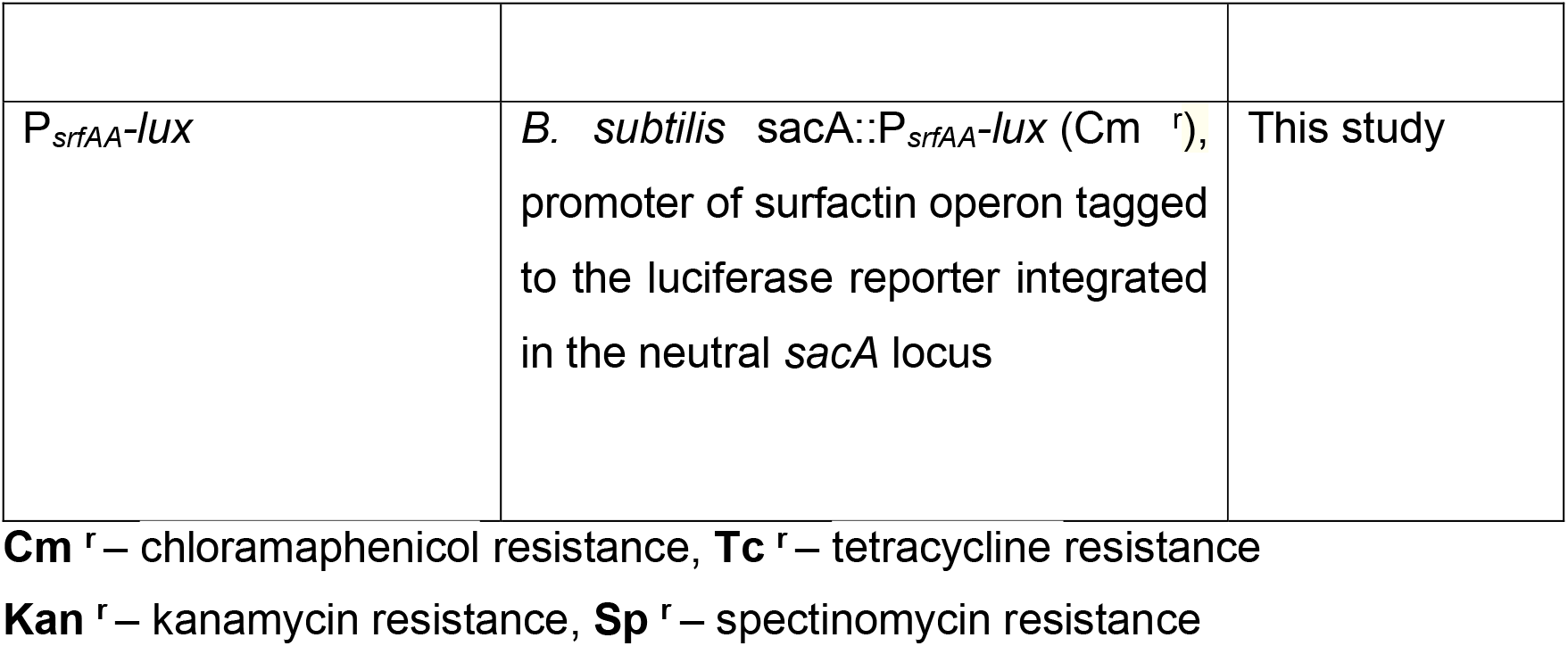
**List of strains used in the study**

| **Strain** | **Description** | **Source or reference** |
| --- | --- | --- |
| *B. subtilis* | Wild type | 9 |
| *B. subtilis* PY79 |  | 10 |
| *B. subtilis natto* |  | Rotem Sorek lab, WIS, Israel |
| *B. atrophaeus* 1942 |  | Rotem Sorek lab, WIS, Israel |
| *B. amyloliquefaciens* *plantarum* FZB42 |  | Rotem Sorek lab, WIS, Israel |
| *B. pumilus* ATCC 70161 |  | Rotem Sorek lab, WIS, Israel |
| *B. megaterium* isolate VS1 |  | Ayelet Fishman Lab, Technion, Israel |
| *B. simplex* SHB26 |  | Lab stock |
| *B. mycoides* AH621 |  | Rotem Sorek lab, WIS, Israel |
| *B. thuringiensis* subspecies *kurstaki* strain HD73 |  | Rotem Sorek lab, WIS, Israel |
| *B. coagulans* ATCC 10545 |  | BGSC 61A2 |
| *B. clausii* *domuvar* |  | BGSC 17A1 |
| $\Delta srfAA$-$srfAD$ ($\Delta srfAA$) | *B. subtilis* $\Delta srfAA::Em^r$ deficient in surfactin synthesis | <sup>11</sup> |
| $\Delta pksC$-$pksR$ ($\Delta pks$) | *B. subtilis* $\Delta pks::Sp^r$ deficient in bacillaene synthesis | <sup>12</sup> |
| $\Delta bacA$-$yfwG$<br>($\Delta bac$) | *B. subtilis* $\Delta bac::Cm^r$ deficient in bacilysin synthesis | This study |
| $\Delta ppsA$-$ppsE$<br>($\Delta pps$) | *B. subtilis* $\Delta pps::Tc^r$ deficient in plipastatin synthesis | This study |
| $\Delta srfAA$ $\Delta pks$ | *B. subtilis* $\Delta srfAA::Em^r$, $\Delta pks::Sp^r$, deficient in surfactin and bacillaene synthesis | This study |
| $\Delta srfAA$ $\Delta bac$ | *B. subtilis* $\Delta srfAA::Em^r$, $\Delta bac::Cm^r$, deficient in surfactin and bacilysin synthesis | This study |
| $\Delta srfAA$ $\Delta pps$ | *B. subtilis* $\Delta srfAA::Em^r$, $\Delta pps::Tc^r$, deficient in surfactin and plipastatin synthesis | This study |
| $\Delta pks$ $\Delta bac$ | *B. subtilis* $\Delta pks::Sp^r$, $\Delta bac::Cm^r$, deficient in bacillaene and bacilysin synthesis | This study |
| $\Delta pks$ $\Delta pps$ | *B. subtilis* $\Delta pks::Sp^r$, $\Delta pps::Tc^r$, deficient in bacillaene and plipastatin synthesis | This study |
| $\Delta bac$ $\Delta pps$ | *B. subtilis* $\Delta bac::Cm^r$, $\Delta pps::Tc^r$, deficient in bacilysin and plipastatin synthesis | This study |
| $\Delta srfAA \Delta pks$<br>$\Delta bac \Delta pps$ ($\Delta 4$) | *B. subtilis* $\Delta srfAA::Em^r$, $\Delta pks::Sp^r$, $\Delta bac::Cm^r$, $\Delta pps::Tc^r$, deficient in surfactin, bacillaene, bacilyisin and plipastatin synthesis | This study |
| $\Delta murP$ | DNA was extracted from *B. subtilis* 168 $\Delta murP$, and transferred into *B. subtilis* 3610 as described in materials and methods. $Kan^r$ | 13 |
| $\Delta nagP$ | DNA was extracted from *B. subtilis* 168 $\Delta nagP$, and transferred into *B. subtilis* 3610 as described in materials and methods. $Kan^r$ | 13 |
| $\Delta prkC$ | DNA was extracted from *B. subtilis* 168 $\Delta prkC$, and transferred into *B. subtilis* 3610 as described in materials and methods. $Kan^r$ | 13 |
| $\Delta ykoG$ | DNA was extracted from *B. subtilis* 168 $\Delta ykoG$, and transferred into *B. subtilis* 3610 as described in materials and methods. $Kan^r$ | 13 |
| $\Delta comA$ | DNA was extracted from *B. subtilis* 168 $\Delta comA$, and transferred into *B. subtilis* 3610 as described in materials and methods. $Kan^r$ | 13 |
| $\Delta ywqM$ | *B. subtilis* $\Delta ywqM::Tc^r$ | This study |
| $\Delta ykoG \Delta ywqM$ | *B. subtilis* $\Delta ywqM::Tc^r \Delta ykoG::kan^r$ | This study |
| $\Delta codY$ | *B. subtilis* $\Delta codY::Kan^r$ | 14 |
| $\Delta spo0A$ | *B. subtilis* $\Delta spo0A::Sp^r$ | 14 |
| $\Delta degU$ | *B. subtilis* $\Delta degU::Tc^r$ | 15 |
| $P_{srfAA-yfp}$ | *B. subtilis* *sacA::P<sub>srfAA-yfp</sub>* (Sp<sup>r</sup>), promoter of surfactin operon tagged to the YFP reporter integrated in the neutral *amyE* locus | Avigdor Eldar Lab, TAU, Israel |
| $P_{pksC-mKate}$ | *B. subtilis* *pyrD::P<sub>pksC-mKate</sub>* (Cm<sup>r</sup>), promoter of bacillaene operon tagged to the mKate reporter integrated in the neutral *pyrD* locus | This study |
| $P_{bacA-gfp}$ | *B. subtilis* *amyE::P<sub>bacA-gfp</sub>* (Cm<sup>r</sup>), promoter of bacilysin operon tagged to the GFP reporter integrated in the neutral *amyE* locus | This study |
| $P_{ppsA-gfp}$ | *B. subtilis* *amyE::P<sub>ppsA-gfp</sub>* (Cm<sup>r</sup>), promoter of plipstatin operon tagged to the GFP reporter integrated in the neutral *amyE* locus | This study |
| $P_{srfAA-yfp}, P_{pksC-mKate}$ | *B. subtilis* *sacA::P<sub>srfAA-yfp</sub>* (Sp<sup>r</sup>), *pyrD::P<sub>pksC-mKate</sub>* (Cm<sup>r</sup>), double reporter to monitor expression of surfactin and bacillaene | This study |
| $P_{pksC-lux}$ | *B. subtilis* *sacA::P<sub>pksC-lux</sub>* (Cm<sup>r</sup>), promoter of bacillaene operon tagged to the luciferase reporter integrated in the neutral *SacA* locus | 14 |
| $P_{srfAA-lux}$ | *B. subtilis* *sacA::P<sub>srfAA-lux</sub>* (Cm<sup>r</sup>),<br>promoter of surfactin operon tagged<br>to the luciferase reporter integrated<br>in the neutral *sacA* locus | This study |

**Cm<sup>r</sup>** – chloramphenicol resistance, **Tc<sup>r</sup>** – tetracycline resistance
**Kan<sup>r</sup>** – kanamycin resistance, **Sp<sup>r</sup>** – spectinomycin resistance

**Supplementary Table S2.**
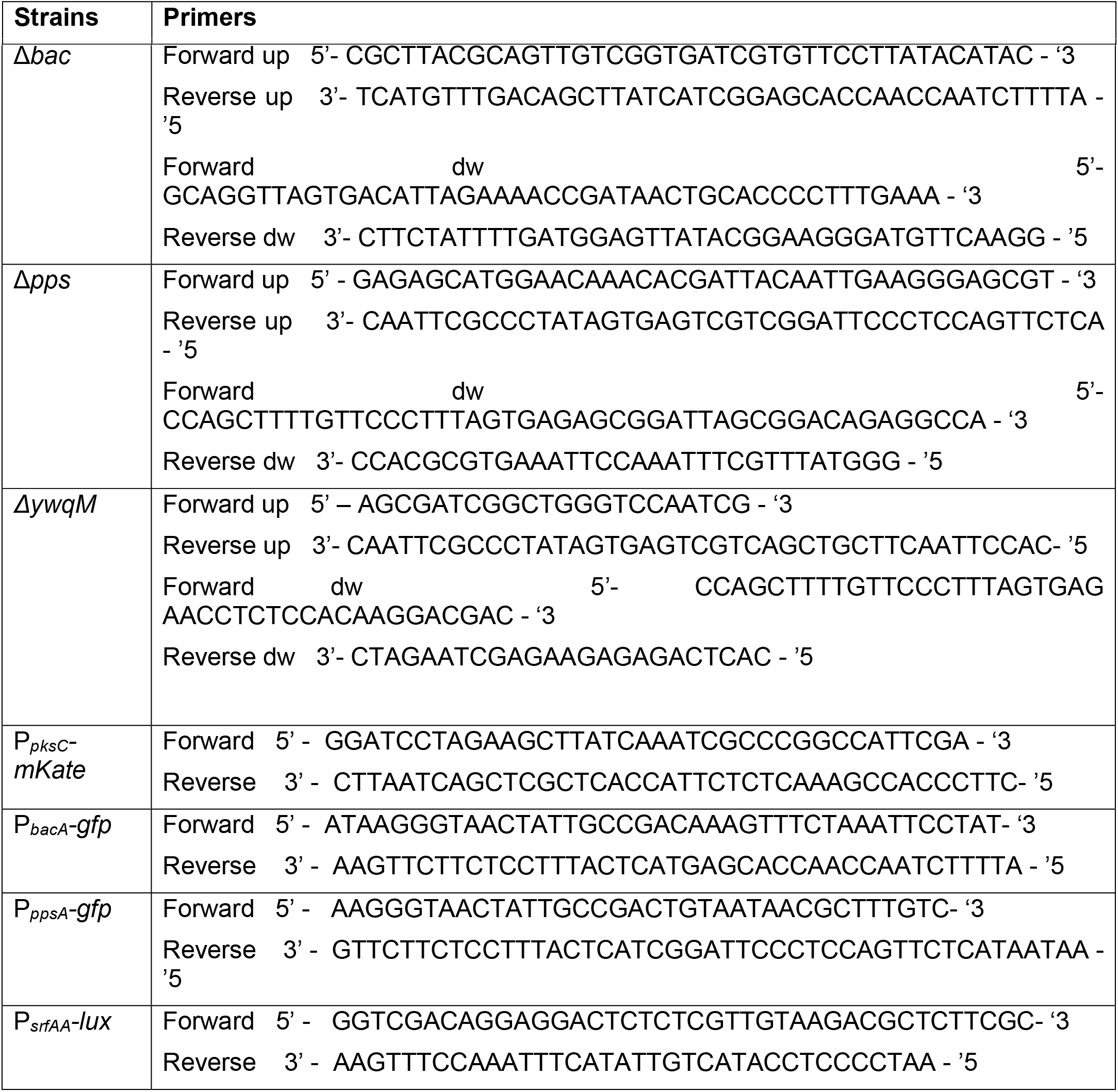
**Primes used in the study**

**Supplementary Table S3.**
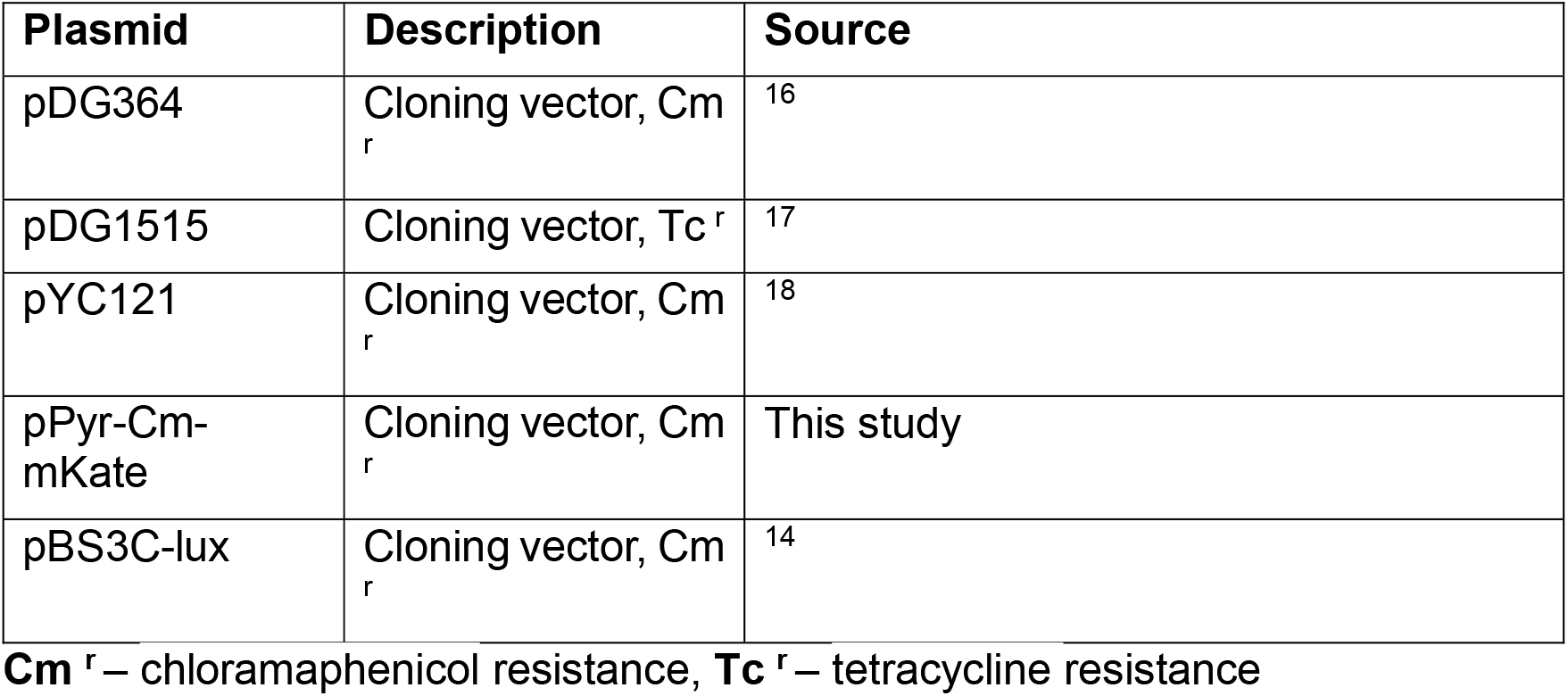
**Plasmids used in the study**

Plasmids used in the study
| Plasmid | Description | Source |
| --- | --- | --- |
| pDG364 | Cloning vector, Cm<sup>r</sup> | <sup>16</sup> |
| pDG1515 | Cloning vector, Tc<sup>r</sup> | <sup>17</sup> |
| pYC121 | Cloning vector, Cm<sup>r</sup> | <sup>18</sup> |
| pPyr-Cm-mKate | Cloning vector, Cm<sup>r</sup> | This study |
| pBS3C-lux | Cloning vector, Cm<sup>r</sup> | <sup>14</sup> |

**Cm<sup>r</sup>** – chloramphenicol resistance, **Tc<sup>r</sup>** – tetracycline resistance

**Table S4:** Transcriptional reporter analyis. A full analysis of the fluorescence of the indicated reporter strains grown as in Figure 4A

|  | **P<sub>pksC</sub> - mKate** |  | **P<sub>srfAA</sub> - yfp** |  | **P<sub>bacA</sub> - gfp** |  | **P<sub>ppsA</sub> - gfp** |  |
| --- | --- | --- | --- | --- | --- | --- | --- | --- |
|  | mKate % | mKate mean | YFP % | YFP mean | GFP % | GFP mean | GFP % | GFP mean |
| **NC** | 18.63 | 1457.33 | 11.63 | 1308.00 | 12.77 | 1449.33 | 0.90 | 1338.67 |
| **vs *B. subtilis*** | 17.87 | 1417.00 | 10.93 | 1261.00 | 12.53 | 1455.33 | 1.03 | 1321.00 |
| **vs *B. natto*** | 17.43 | 1423.67 | 11.47 | 1275.33 | 11.73 | 1453.67 | 0.97 | 1391.00 |
| **vs *B. atrophaeus*** | 14.73 | 1395.00 | 9.33 | 1253.67 | 8.60 | 1400.00 | 0.70 | 1566.33 |
| **vs *B. amyloliquefaciens*** | 16.63 | 1409.00 | 9.03 | 1248.00 | 9.27 | 1408.67 | 0.77 | 1541.67 |
| **vs *B. pumilus*** | 28.97 | 1662.33 | 15.13 | 1434.67 | 14.57 | 1498.67 | 2.63 | 1482.67 |
| **vs *B. megaterium*** | 30.80 | 1772.00 | 17.93 | 1505.33 | 16.33 | 1548.00 | 3.43 | 1427.33 |
| **vs *B. simplex*** | 27.60 | 1696.67 | 15.43 | 1450.67 | 17.27 | 1565.00 | 3.40 | 1533.67 |
| **vs *B. Mycoides*** | 31.43 | 1781.67 | 18.67 | 1509.67 | 17.57 | 1633.33 | 3.53 | 1536.00 |
| **vs *B. thuringiensis*** | 31.37 | 1770.00 | 18.83 | 1511.33 | 18.30 | 1677.00 | 3.50 | 1461.67 |
| **vs *B. coagulans*** | 28.83 | 1743.00 | 17.37 | 1495.67 | 17.30 | 1575.33 | 3.23 | 1456.00 |
| **vs *B. clausii*** | 28.47 | 1740.00 | 17.13 | 1493.00 | 16.20 | 1558.33 | 2.73 | 1478.67 |

## Supporting Methods

### Strain construction

All deletion mutants were derived from *B. subtilis* NCIB 3610 WT. Deletions mutants were constructed using standard methods as described in ^1,2^. For polymerase chain reactions, primers and plasmids used in this study are listed in supplementary Table S2 and S3 respectively. Briefly genomic regions of around 1000 bp upstream and downstream of NRP operon were amplified from *B. subtilis* chromosomal DNA. Deletions were generated using the long-flanking homology (LFH) PCR mutagenesis protocol of ^3^, replacing an endogenous locus with a resistance gene from either pDG1515 (Δ*pps* and Δ*ywqM*) and pDG364 (Δ*bac*). DNA was extracted with the Wizard Genomic DNA Purification Kit (Promega) transformed into strain *B. subtilis* PY79. Genomic DNA of the recipient was transformed into *B. subtilis* NCIB 3610 or deletion mutants as described ^4^. All mutant strains were confirmed for accurate integration by PCR. Restriction enzymes and Phusion HF DNA Polymerase were purchased from New England BioLabs.

All cloning experiments were performed with *B. subtilis* NCIB 3610 and *E*. *coli* DH5α and by restriction free cloning ^5^. pYC121 plasmid was used as template for P*_bacA_*-*gfp* (bacilysin) and P*_ppsA_*-*gfp* (plipastatin) and plasmid pPyr-Cm – mKate for P*_pksC_*-*mKate*. Briefly, PCR fragments of promoter regions were amplified from *B. subtilis* chromosomal DNA, using primers listed in supplementary Table S2 and ligated into their respective plasmids (Table S3). The ligated plasmids were then transformed into *E*. *coli* DH5α. Positive reporters were inserted by using double homologous recombination into neutral integration sites *(amyE)* and *(pyrD)* in the genome of *B. subtilis* by inducing natural competence ^4^ and confirmed by PCR. Selective media for cloning purposes were prepared with LB broth or LB-agar. *B. subtilis* P*_srfAA_*-*yfp* was a kind gift from the lab of Avigdor Eldar, TAU Israel.

To create double-labelled strain of *B. subtilis* P*_pksC_*-*mKate*, P*_srfAA_*-*yfp*, genomic DNA from *B. subtilis* P*_srfAA_*-*yfp* was integrated into neutral integration sites (*sacA*) in the genome of *B. subtilis* P*_pksC_*-*mKate*. Dually labelled strains were confirmed for successful transformation by PCR. sSelective media for cloning purposes were prepared with LB broth or LB-agar using antibiotics at the following final concentrations: 10 μg/ml chloramphenicol (Amersco) and 10 μg/ml spectinomycin (Tivan biotech).

For the generation of the Luminescence reporter P*_srfAA_*-*lux* we performed restriction free cloning ^5^ using the plasmid pBS3C-lux ^6^, which contains a functional luciferase operon *(luxABCDE)*. Briefly, promoter region was amplified from *B. subtilis* chromosomal DNA, using primers listed in supplementary Table S2 and ligated into plasmid. The ligated plasmids were then transformed into *E*. *coli* DH5α. Positive reporters were inserted by using double homologous recombination into neutral integration sites *(sacA)* in the genome of *B. subtilis* by inducing natural competence ^4^. Selective media for cloning purposes were prepared with LB broth or LB-agar using antibiotics at the following final concentrations: 10 μg/ml chloramphenicol (Amersco).

### Constructing ofs pPyr-Cm - mKate plasmid

Primer pairs for generating DNA fragments with varying overlapping ends by PCR were designed using SnapGene Viewer program. Briefly, mKate gene was first amplified from pDR183 using primers mKate forward - 5’-GGATCCTAGAAGCTTATCAAATCGCCCGGCCATTCGA - ‘3, mKate reverse 3’- CTGTCAAACATGAGAATTCGTCATCTGTGCCCCAGTTTGC - ‘5, pPyr-Cm backbone was amplified using primers vector forward 5’-GCAAACTGGGGCACAGATGACGAATTCTCATGTTTGACAG - ‘3, vector reverse 3’- TAATCAGCTCGCTCACCATATAAGCTTCTAGGATCCTGAG - ‘5. The PCR products were then assembled using NEBuilder^®^ HiFi DNA Assembly Master Mix, based on manufacture protocol and the reaction mix was incubated for 50°C. Following incubation, the Gibson assembly products were then treated with DpnI digestion the transferred into competent *E*. *coli* DH5α cells. Positive reporters were selected and the insertion of mKate was confirmed by PCR. Selective media for cloning purposes were prepared with LB broth or LB-agar using antibiotics 10 μg/ml chloramphenicol (Amersco).

### Extraction of DNA, EPS and PG

DNA was extracted using Wizard^®^ Genomic DNA Purification Kit. Spectrophotometric analysis was performed using NanoDrop spectrophotometer (DNA ≈ 100ng/μL) and DNA was stored at −80 °C for further use.

EPS was extracted from the bacilli biofilms grown at 48 h at 30°C in B4 medium. Biofilm colonies were scrapped and suspended in phosphate-buffered saline (137 mM NaCl, 2.7 mM KCl, 10 mM Na2HPO4, 1.8 mM KH_2_PO_4_), were mildly sonicated, and were then centrifuged to remove the cells. The supernatant was collected and mixed with five volumes of ice-cold isopropanol and incubated overnight at 4°C. Samples were then centrifuged at 7,500 × g for 10 min at 4°C. Pellets were suspended in a digestion mix of 0.1 M MgCl_2_, 0.1 mg/ml of DNase, and 0.1 mg/ml of RNase, and were incubated for 4 h at 37°C. Samples were extracted twice with phenol-chloroform. The aquatic fraction was dialyzed for 48 h with Slide-A-Lyzer dialysis cassettes by Thermo Fisher, 3,500 molecular weight cut-off, against distilled (d)H_2_O. Samples were lyophilized and lyophilized fraction was dissolved in 500 μl of dH_2_O. The fractions were stored at −80 °C for further use.

Peptidoglycan (PG) was extracted from bacilli as previously described in ^7,8^. Briefly, bacilli were grown in volume three hundred milliliter of B4 medium to an OD600 ~1.2. Cells were collected by centrifugation, washed with 0.8% NaCl, resuspended in hot 4% SDS, boiled for 30 min, and incubated at room temperature (RT) overnight. The suspension was then centrifuged to collect the pellet, and washed five times with dH_2_O to remove SDS. The pellet was then suspended in 25 mL 100 mM Tris–HCl, pH 7.5. 25 μL of RNAse solution (10 mg/mL), 25 μL of DNAse solution (10 mg/mL), and 250 μL of 1 M MgSO_4_ and incubated for 4h at 37 °C, with gentle shaking. This was followed by treatment with 25 μL of trypsin (10 mg/mL) and 250 μL of 1 M CaCl2 and incubated at 37 °C for 16 h, with gentoile shaking. The insoluble material was then centrifuged at 6000 × *g* for 10 min at room temperature, wash once with water. The material was again resuspended in 4% SDS, boiled for 30 min, and incubated at room temperature (RT) overnight. The material was then washed five times with washed five times with dH_2_O to remove SDS. Pellet containing PG was dissolved in dH_2_O, was stored at −20 °C for further use.

